# Prospective modelling of operational offshore windfarms on the distribution of marine megafauna in the southern North Sea

**DOI:** 10.1101/2020.12.16.423009

**Authors:** Auriane Virgili, Sophie Laran, Matthieu Authier, Ghislain Doremus, Olivier Van Canneyt, Jérôme Spitz

**Affiliations:** Observatoire Pelagis, UMS 3462 CNRS - La Rochelle Université, La Rochelle, France; Centre d’Etudes Biologiques de Chizé (CEBC), UMR 7372 du CNRS-La Rochelle Université, 79360 Villiers-en-Bois, France

**Keywords:** BAG design, Counterfactuals, Distribution and abundance, Prospective impact

## Abstract

Intense development of Offshore Wind Farms (OFWs) has occurred in the North Sea with several more farms planned for the near future. These OFWs pose a threat to marine megafauna stressing the need to mitigate the impact of human activities. To help mitigating impacts, the Before After Gradient (BAG) design was proposed. We thus explored the use of the BAG method on megafauna sightings recorded at different distances from wind farms in the southern North Sea. We predicted intra-annual variability in species distribution, then correlated species distribution with the presence of operational OFWs and investigated the potential impact the operation of prospective OFWs may have on species distribution. Three patterns of intra-annual variability were predicted: species present most abundantly in spring, in winter or all year-round. We recommend that future OFW constructions be planned in summer and early fall to minimise impact on cetaceans and that offshore areas off northern France and Belgium be avoided to minimise impact on seabirds. Our prospective analysis predicted an increased or a decreased density with the operation of prospective OFWs. Prospective approaches, using e.g. a BAG design, are paramount to inform species conservation as they can forecast the likely responses of megafauna to anthropogenic disturbances.

## 1. Introduction

Anthropogenic activity in the southern region of the North Sea is among the most intense in the world. A strongly used shipping lane crosses the Dover Strait and the Belgian and British waters are used for intense fisheries as well as for offshore wind farm construction (Halpern et al., 2008). There has been a marked increase in marine renewable energy projects in the southern North Sea in recent years, particularly in offshore wind farms (OFWs), the number of which is expected to more than double in the coming years (from currently six to ± 14; https://www.4coffshore.com/offshorewind/). The development of these OFWs pose various threats to marine megafauna (*i.e*. marine mammals, seabirds, large fish and marine turtles) living in and travelling through the area, including: (1) direct mortalities due to collisions between seabirds and wind turbines (Drewitt et al., 2006); (2) physiological stress, such as lesions and hearing loss due to pile driving (Board & National Research Council, 2005; Tougaard et al., 2005; Thomsen et al., 2006; Dähne et al., 2013); and (3) disturbances due to pile driving and intense marine traffic, which can induce changes in distribution, behaviour, reproduction or foraging activity patterns, *e.g*. by displacing individuals from suitable habitats (Drewitt et al., 2006; Brandt et al., 2011; Bergström et al., 2014; Vallejo et al., 2018; Mendel et al., 2019). Harbour porpoises (*Phocoena phocoena*) and various seabird species (*e.g*. northern gannets *Morus bassanus*, northern fulmars *Fulmarus glacialis*) occur in high densities in the southern part of the North Sea (Lambert et al., 2017) and are therefore significantly at risk of being affected by the spread of wind farms. It is important that we understand the threats that anthropogenic developments and activities pose to marine biodiversity to allow development of specific recommendations to mitigate them. This is the aim of the EU’s Directive 2014/89/EU, which calls for efficient marine spatial planning in European waters (Douvere, 2008; European Parliament and Council Directive 2014/89/EU).

The mitigation of threats to biodiversity comprises four sequential steps, based on the mitigation hierarchy, namely: (1) avoidance, (2) minimisation, (3) restoration and (4) offsetting (Kiesecker et al., 2010). Avoidance involves taking measures to prevent impacts (*e.g*. excluding breeding areas in prospective development plans). Minimisation measures reduce the duration, intensity and extent of impacts that cannot be avoided (*e.g*. making technological adjustments to reduce noise). When impacts cannot be completely avoided or minimised, restoration efforts aim to return an ecosystem to a pre-impact state. After the implementation of the first three steps, any residual impacts can be offset - or compensated for - in order to balance the detrimental effects with positive effects (*e.g*. preventing future habitat degradation; Kiesecker et al., 2010).

Avoidance is the easiest and most effective step, particularly in the context of wind farming. It requires that the initial state of the biodiversity in the prospected area be thoroughly investigated in order to assess when the construction phase would have the least impact on species and where the wind farm should be established to minimise impacts during the operational phase. This assessment critically hinges on accurate knowledge of species distributions, particularly on a seasonal and monthly scale (Foley et al., 2010; Santos et al., 2019). Knowledge of the monthly distribution of marine megafauna (marine mammals and seabirds in this particular area) is essential in a marine spatial planning context because these animals are highly mobile and their distributions change depending on their life cycle. Animals are particularly sensitive to disturbance during certain life stages, *e.g*. during mating or breeding seasons, therefore accurate seasonal and monthly distribution patterns are necessary in order to reduce the spatial and/or temporal overlap between any disruptive activities and the sensitive stages or with key aggregation areas of the species (Gilles et al., 2011; Whitt et al., 2013; Pitchford et al., 2016).

The Before After Control Impact (BACI) method (Green, 1979), consisting in comparing an impact location (*e.g*. a wind farm) to a control location (unaffected by humans) before and after the human intervention, is often used to meet avoidance objectives and to study the effect of offshore wind farm on species (Bergström et al., 2013; Degraer et al., 2018; Wilber et al., 2018). However, this method has limitations (*e.g*. difficulty in finding a suitable control location, assumption that all wind farms are equivalent) and Methratta (2020) has proposed to integrate an alternative approach into fisheries surveys, the Before After Gradient (BAG) design, to overcome these limitations. Sampling is carried out along a spatial gradient at increasing distances from impact locations before and after the construction, the distance to wind farms therefore plays a very important role. With data collected with the BAG design, Methratta (2020) proposed to perform statistical analysis, such as generalised linear or additive models, using the distance to wind farms as main explanatory variable. Relationships between samples and distance to wind farms are thus obtained before and after human intervention. The BAG design for fisheries resources is very promising and can also be applied to animals that depend on this resource such as top predators. In our study, we therefore explored the use of the BAG method on seabird and marine mammal sightings recorded at different distances from two wind farms in the southern North Sea.

We conducted six aerial surveys in the southern North Sea (from 50.75 to 52°N and from 1 to 3.5°E) between April 2017 and May 2018, allowing regular monitoring over a whole year so that we could study the intra-annual variability in the distribution and abundance of marine megafauna. According to the BAG framework, we used density surface models to (1) predict the monthly distribution and abundance of species (Guisan & Thuiller, 2005) and (2) to correlate species distributions with the distance to two operational OFWs in the area. Based on this empirical correlation, we (3) investigated the potential impact the operation of prospective OFWs may have on species distribution, on a monthly basis. These prospective OFWs are projects currently in concept or authorised for construction but for which construction has not yet begun (https://www.4coffshore.com/offshorewind/). As suggested by Methratta (2020), our aim was to use the distance to operational OFWs as predictor in the models to establish relationships and then use these relationships to assess how species distribution would change if OFWs currently approved were implemented and fully operational in the area. We did not assess scenarios of noise or impact reduction during the construction phase of an OFW (Interim Population Consequences of Disturbance - iPCoD model; Harwood & King, 2014; Degraer et al., 2018) nor assess the cumulative noise impact of the construction of multiple OFWs (Disturbance Effects of Noise on the Harbour Porpoise Population in the North Sea - DEPONS model, Nabe-Nielsen et al., 2018).

A good understanding of the current monthly species distributions and the potential impacts of the increasing wind farm activity in the future will allow for clear guidelines and recommendations defining when and where wind farm construction would have the least harmful impact ecologically and will facilitate the effective mitigation and management of marine activities in the area.

## 2. Materials and Methods

### 2.1. Species of interest

We focused on six species or species groups in this study, namely: the harbour porpoise (*Phocoena phocoena*), auks (*Uria aalge* and *Alca torda*), the black-legged kittiwake (*Rissa tridactyla*), cormorants (*Phalacrocorax carbo* and *Phalacrocorax aristotelis*), the northern fulmar (*Fulmarus glacialis*) and the northern gannet (*Morus bassanus;* Table 1). Only harbour porpoises and the seabirds listed above were present in numbers adequate for analysis. Harbour porpoises, northern gannets and northern fulmars could be clearly identified from the aircraft and analyses could be conducted at the species level. The other species needed to be considered as groups (cormorants, auks, black-legged kittiwakes; Table 1) based on taxonomic and morphological criteria.

**Table 1.** Composition of species groups.

| Family | Group names | Group composition |  |
| --- | --- | --- | --- |
| **Alcidae** | Auks | Common guillemots<br>Razorbills | *Uria aalge*<br>*Alca torda* |
| **Laridae** | Black-legged kittiwakes | Black-legged kittiwakes | *Rissa tridactyla* |
|  |  | Indeterminate small gulls | *Rissa tridactyla* /<br>*Chroicocephalus ridibundus* /<br>*Ichtyaetus melanocephalus* |
| **Phalacrocoracidae** | Cormorants | Great cormorants<br>European shags | *Phalacrocorax carbo*<br>*Phalacrocorax aristotelis* |
| **Phocoenidae** | Harbour porpoise |  | *Phocoena phocoena* |
| **Procellariidae** | Northern fulmar |  | *Fulmarus glacialis* |
| **Sulidae** | Northern gannet |  | *Morus bassanus* |

The black-legged kittiwake group also incorporates sightings of small gulls that could not be confidently identified to species level by observers (~ 13% of sightings included in the kittiwake group). Based on the geographic area, unidentified gulls could have been black-legged kittiwakes, black-headed gulls (*Chroicocephalus ridibundus*) or Mediterranean gulls (*Ichtyaetus melanocephalus*), however, these two gull species represented a very small proportion of the observed gulls that could be clearly identified (less than 30 sightings out of a total of 1613 gulls recorded in all flight sessions). The unidentified gull sightings also occurred offshore, unlike the identified black-headed and Mediterranean gulls, which were recorded in-shore, so it is likely that many of these unidentified sightings were in fact kittiwakes. We therefore do not expect the inclusion of unidentified gulls to bias results.

### 2.2. Data collection and data processing

The study area covered about 9,400 km^2^ and included the Belgian waters and parts of the French and English waters in the North Sea (from 50.75 to 52°N and from 1 to 3.5°E; Figure 1). Aerial visual surveys were conducted in 2017 on April 6-7, June 13-14, August 7-8, and December 4-5, and in 2018 on March 5-7 and May, 4-5. Aerial transects are consistent with a BAG design with sampling carried out at different distances from two wind farms, operational since 2009 and 2010, the Thanet wind farm, off the east end of Kent (England) and the Thorntonbank wind farm between the Belgian and Dutch waters (Figure 1).

**Figure 1.**
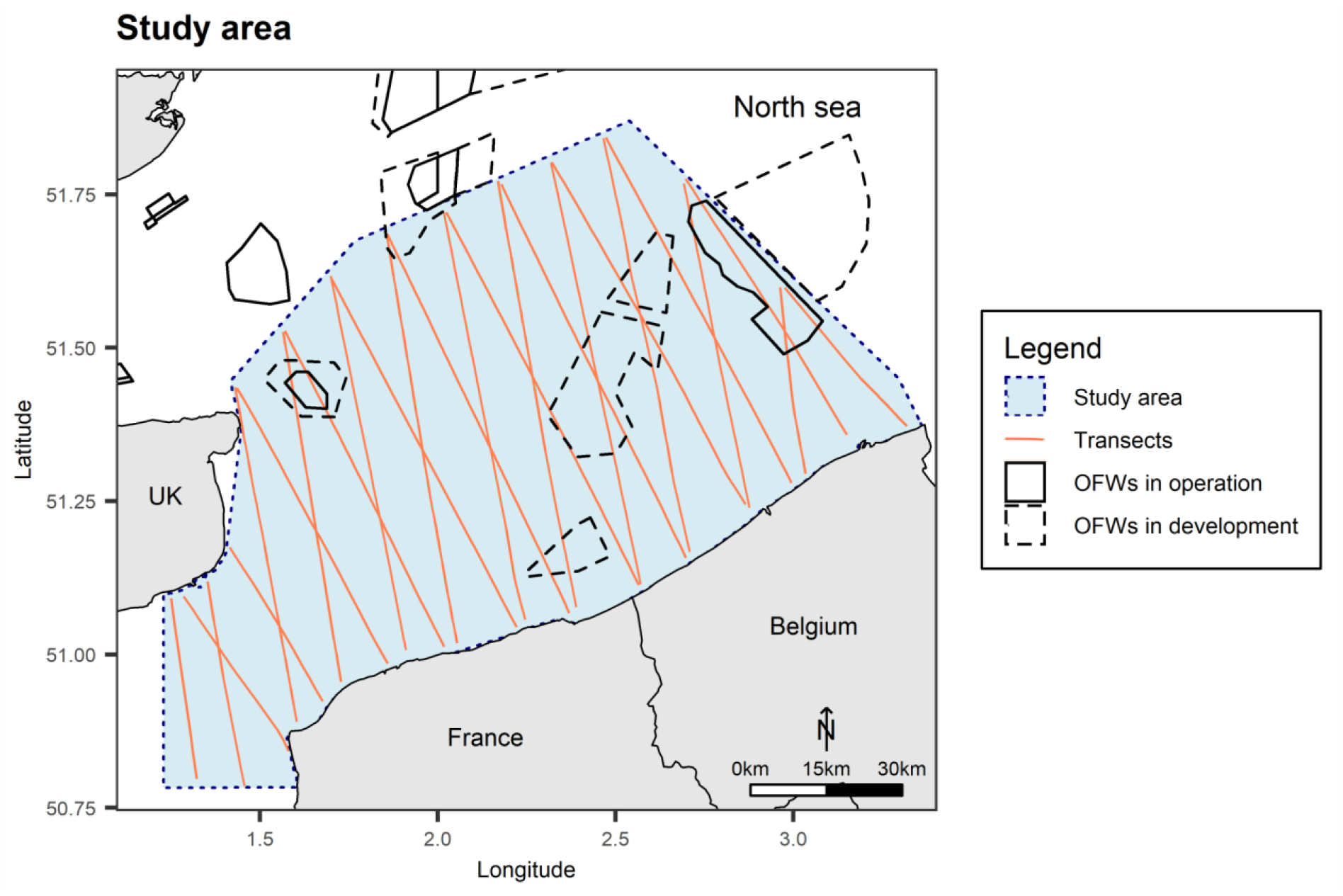
Study area and sampling plan. The dotted blue box delineates the study area, solid orange lines represent the sampling plan (for monthly detail refer to Appendix A), solid black boxes represent the OFWs in operation and dashed black boxes the OFWs in development (OFWs: Offshore Wind Farms; UK: United Kingdom).

We used a multi-target protocol (described in details in Lambert et al., 2019) to simultaneously record harbour porpoise and seabird sightings. Sightings of seabirds were recorded within a 200 m strip on each side of the aircraft (strip transect protocol), while for harbour porpoises, the perpendicular distance to the transect was recorded for distance sampling analyses (line transect protocol; Buckland et al., 2001). Linear transects were designed using the equal spaced zig-zag option of the software ‘Distance’ version 6.2 (Buckland et al., 2001; Thomas et al., 2010). This design can produce an unequal coverage probability when study regions are non-rectangular (Strindberg & Buckland, 2004). This is why we used density surface modelling, a model-based approach that relaxes the equal sampling probability assumption of conventional distance sampling by incorporating spatially explicit covariates to obtain a fine-grained density surface in the study area (Miller et al. 2013; Appendix B compares porpoise densities estimated by conventional distance sampling and density surface modelling). Sightings and observation conditions were recorded using ‘SAMMOA 1.0.4’ software (SAMMOA 1.0.4.; http://www.observatoire-pelagis.cnrs.fr/publications/les-outils/article/logiciel-sammoa) at the beginning of each transect and upon any change in weather, flight route, sea state, etc.

Datasets were prepared for analyses using the ‘Marine Geospatial Ecology Tools’ package for ‘ArcGIS 10.3’ (ESRI, 2016; Roberts et al., 2010). Effort data were linearized and segmented into 2.5 km segments of homogenous observation conditions. Only segments recorded with good observation conditions were kept for analysis (*i.e*. Beaufort sea-state between 0 and 3 and subjective observation conditions from medium to excellent).

### 2.3. Density surface modelling

Density surface models (DSM; Miller et al., 2013) allow us to investigate the relationships between species and their environment and predict their distributions. To do this we included both static and dynamic environmental variables, considered as potential drivers of prey availability.

#### 2.3.1. Environmental variables

Environmental variables are described by two subsets, the static and dynamic variables (Table 2). Static variables included distance to the coast, the minimum Euclidian distance to the OFWs, and for seabirds, the distance to the closest colony. Two types of OFWs were included in the analysis: 1) operational OFWs, from which the effect on species distributions were quantified; and 2) OFWs in development – planned farms for which construction had not yet begun (https://www.4coffshore.com/offshorewind/) - which were used for the prospective analysis. Colonies were mapped for each seabird group (Legroux et al., 2017; Appendix C) and the distance between the centre of each segment and the nearest colony was calculated. We also included a residual and time-varying spatial effect in the models to account for spatial autocorrelation (Wikle & Royle, 2002; see supplementary material in Appendix D for details). This spatial effect can be interpreted as the effect of environmental covariates not included in the models, but implies that predictions are restricted to the study area.

**Table 2.** Candidate environmental variables used for the density surface modelling. **A**: The distance from sighting location to the coast was calculated in ArcGIS 10.3. **B**: The distance from sighting location to the operational wind farms was calculated in ArcGIS 10.3 using the wind farm perimeters provided by http://www.marineatlas.be/en/data for Belgium and by https://www.thecrownestate.co.uk/en-gb/resources/maps-and-gis-data/ for the United Kingdom. **C**: The distance to the wind farms in development was calculated in ArcGIS 10.3 using the wind farm perimeters provided by https://www.4coffshore.com/offshorewind/. **D**: The distance to the colonies was calculated in ArcGIS 10.3 from the location of the colonies (Appendix C). **E:** Mean sea surface temperatures were extracted from NASA data (https://podaac.jpl.nasa.gov/dataset/JPL_OUROCEAN-L4UHfnd-GLOB-G1SST). **F**: Current speeds were extracted from IFREMER’s MARS 2D model (https://marc.ifremer.fr/resultats/). **G**: Chlorophyll *a* concentrations were extracted from IFREMER’s ECOMARS 3D model (https://marc.ifremer.fr/resultats/). Sea surface temperatures were obtained from satellite data while other variables were derived from predictions of oceanographic models. These models were particularly useful in the case of chlorophyll *a* concentration because satellite data were unavailable in winter due to cloud coverage.

| Environmental variables | Original resolutions | Sources | Justification |
| --- | --- | --- | --- |
| **Physiographic** |  |  |  |
| Distance to coast (km) | - | A | Environment constraint |
| Distance operational to wind farms (km) | - | B | Expected effect of the presence of wind farms: avoidance, attraction |
| Distance to wind farms in development (km) | - | C | Expected effect of the presence of wind farms: avoidance, attraction |
| Distance to colony (km) | - | D | Reproductive constraint |
| **Oceanographic** |  |  |  |
| Sea surface temperature (°C) | 0.01° (≈ 0.8 km), day | E | Variability over time and horizontal gradients of SST reveal front locations, potentially associated with prey aggregations or enhanced primary production |
| Current speed (m.s<sup>-1</sup>) | 750 m, hour | F | Intensity of tidal currents |
| Chlorophyll *a* concentration (mg.m<sup>-3</sup>) | 4 km, week | G | Proxy of primary production and therefore of prey abundance and distribution |

For dynamic variables, we used a 28-day temporal resolution (*i.e*. variables were averaged over the 28 days, or 4 weeks, prior the surveyed day) to account for a time lag between an environmental change and the response of marine megafauna to this new condition (Lambert et al., 2017). Environmental variables were averaged over a 2.5 × 2.5 km grid to match the 2.5 km segment length. Dynamic variables included sea surface temperature, maximum current speed and chlorophyll *a* concentration. Sea surface temperature serves as a proxy for the location of fronts in which prey are concentrated. Maximum current speed was included because the English Channel and the southern North Sea are characterised by intense tidal currents.

Chlorophyll *a* concentration serves as an indicator of prey distribution. Appendix E shows the average situation of each environmental covariate.

All environmental variables were standardised (mean-centred and divided by one standard deviation) prior to model fitting and then back-transformed to the original scale for ease of interpretation.

#### 2.3.2. Density surface modelling

To model the relationships between the number of individuals (response variable) and the environment (predictors), we used generalised additive models (GAMs; Hastie & Tibshirani, 1986; Wood, 2006) with a binomial negative distribution to account for data over-dispersion (Gilles et al. 2016; Isaac et al., 2019). GAMs are extensions of generalised linear models (GLMs) that allow for nonlinear relations between the response variable and predictors. It is important to note that the density surface models we built allowed for seasonal effects in both the intercepts and nonlinear relationships with the environment to predict monthly distributions. This effect was assumed to be cyclical: for example, the average density in January 2018 was correlated with the average density in December 2017 and February 2018 but not with the average density in August 2017 or 2018. An interaction with the month of the survey was included in the smoothing functions so that relationships with the environment can vary by month. This allowed to obtain monthly relationships and then predict monthly distributions even if some months were not sampled during the surveys (see Appendix D for a detailed model description).

We fitted a GAM for each species group and used a model selection procedure to select the best model. To avoid collinearity between variables, we first excluded combinations of variables with Pearson correlation coefficients greater than | 0.7 | (Dormann et al., 2013). Fitted models always included the “Distance to operational wind farms “ for harbour porpoises and the “Distance to operational wind farms” and “Distance to colony” for seabirds to estimate the relationship between the number of individuals and the distance to the operational OFWs, and to assess the effect of distance to colonies on seabird distribution. This systematic inclusion of “Distance to operational wind farms” is akin to the BAG design of Methratta (2020). For the other variables, a whole-subset model selection procedure with the Widely Applicable Information Criterion (Vehtari et al., 2017) selected the variables most predictive of species distribution. A maximum of five and six variables was allowed for harbour porpoises and seabirds.

The best predictive model (corresponding to the lowest WAIC) was used to (i) map monthly relative densities on a 2.5 × 2.5 km grid, and (ii) estimate monthly abundances and their associated 80% credible intervals. To limit extrapolation, we only performed predictions within the sampled environmental envelope.

Empirical relationships with the operational OFWs were then used to predict densities and abundances using all OFWs located in the study area - both those that are operational and those under development. These predictions correspond to counterfactuals, quantifying what would be observed, *ceteris paribus*, in the hypothetical scenario where all planned OFWs had already been completed. The selected models do not enable us to predict the effects of OFW construction on marine megafauna, only what the effects of the OFWs would be if they were all operational. We were therefore able to predict new monthly distributions and abundances for each species group by considering the presence of operational and planned OFWs. We compared the observed monthly distributions and abundances with those predicted from the theoretical scenario using the earth mover’s distance (EMD, R-package ‘move’; Kranstauber et al., 2018). The EMD is defined as the minimal amount of work needed for an earth mover to transform one landscape into another, a value of 0 indicating that the distribution maps match perfectly (Kranstauber et al., 2017).

## 3. Results

### 3.1. Survey effort and sighting data

A total of 8,840 km was flown during the six flight sessions (Table 3). Of the total effort 89% was performed in good conditions, *i.e*. with a Beaufort sea-state strictly less than four and subjective observation conditions between medium and excellent.

**Table 3.**
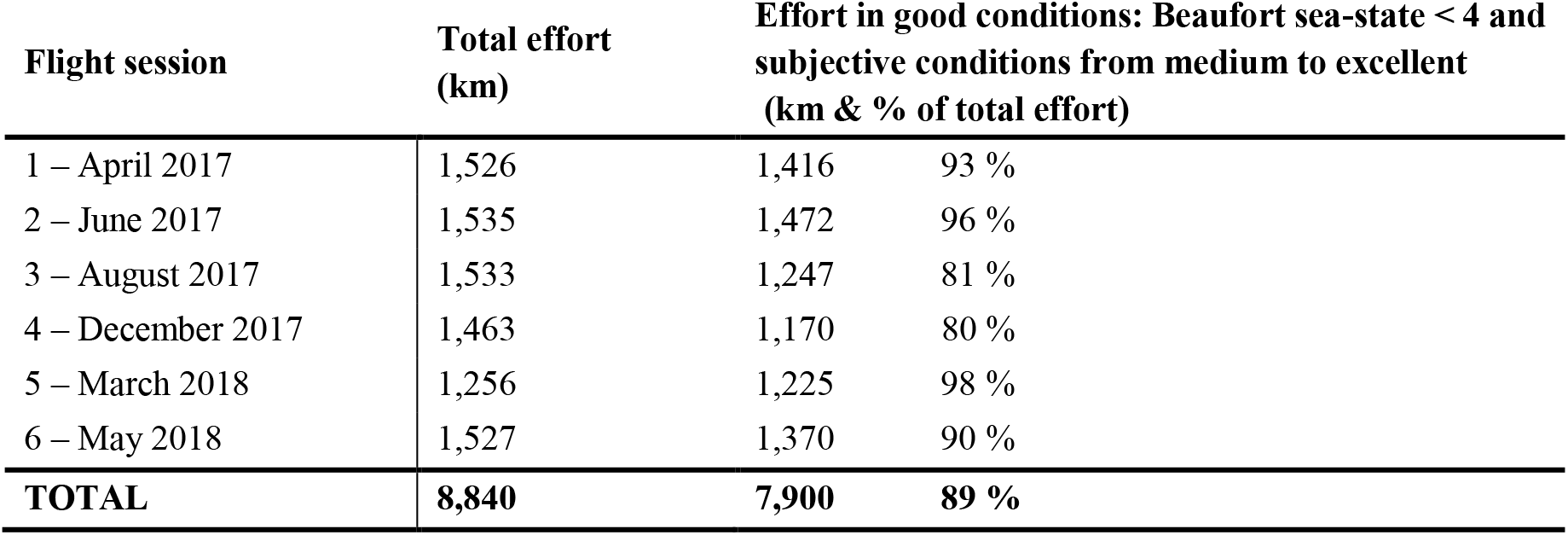
Total effort and effort carried out with optimal observation conditions (in km) per flight session.

| Flight session | Total effort (km) | Effort in good conditions: Beaufort sea-state < 4 and subjective conditions from medium to excellent (km & % of total effort) |  |
| --- | --- | --- | --- |
| 1 – April 2017 | 1,526 | 1,416 | 93 % |
| 2 – June 2017 | 1,535 | 1,472 | 96 % |
| 3 – August 2017 | 1,533 | 1,247 | 81 % |
| 4 – December 2017 | 1,463 | 1,170 | 80 % |
| 5 – March 2018 | 1,256 | 1,225 | 98 % |
| 6 – May 2018 | 1,527 | 1,370 | 90 % |
| **TOTAL** | **8,840** | **7,900** | **89 %** |

Sightings recorded during the six flight sessions, detailed in Appendix F, included 1,120 sightings of harbour porpoises (1,412 individuals) and 6,802 seabird sightings (18,562 individuals). The seabird sightings included 1,218 sightings of auks, 1,592 of black-legged kittiwakes, 134 of cormorants, 186 of northern fulmars and 1,421 of northern gannets. These sightings included both individuals and groups.

### 3.2. Density surface modelling

All species except for cormorants were predicted to be distributed offshore. The predicted distribution of cormorants was very coastal. Three seasonal patterns of abundance could be defined post-hoc: some species were abundant in spring (March to May) and almost absent in summer (June to August); others were abundant in winter (December to February), and the rest showed little seasonal variations throughout the year.

#### 3.2.1. Species abundant in spring and almost absent in summer

##### Harbour porpoises

The highest estimated densities of harbour porpoises were associated with low chlorophyll *a* concentrations, distances to the coast of ± 25-30 km and beyond 50 km, low current speed and low sea surface temperatures (Figure 2).

**Figure 2.**
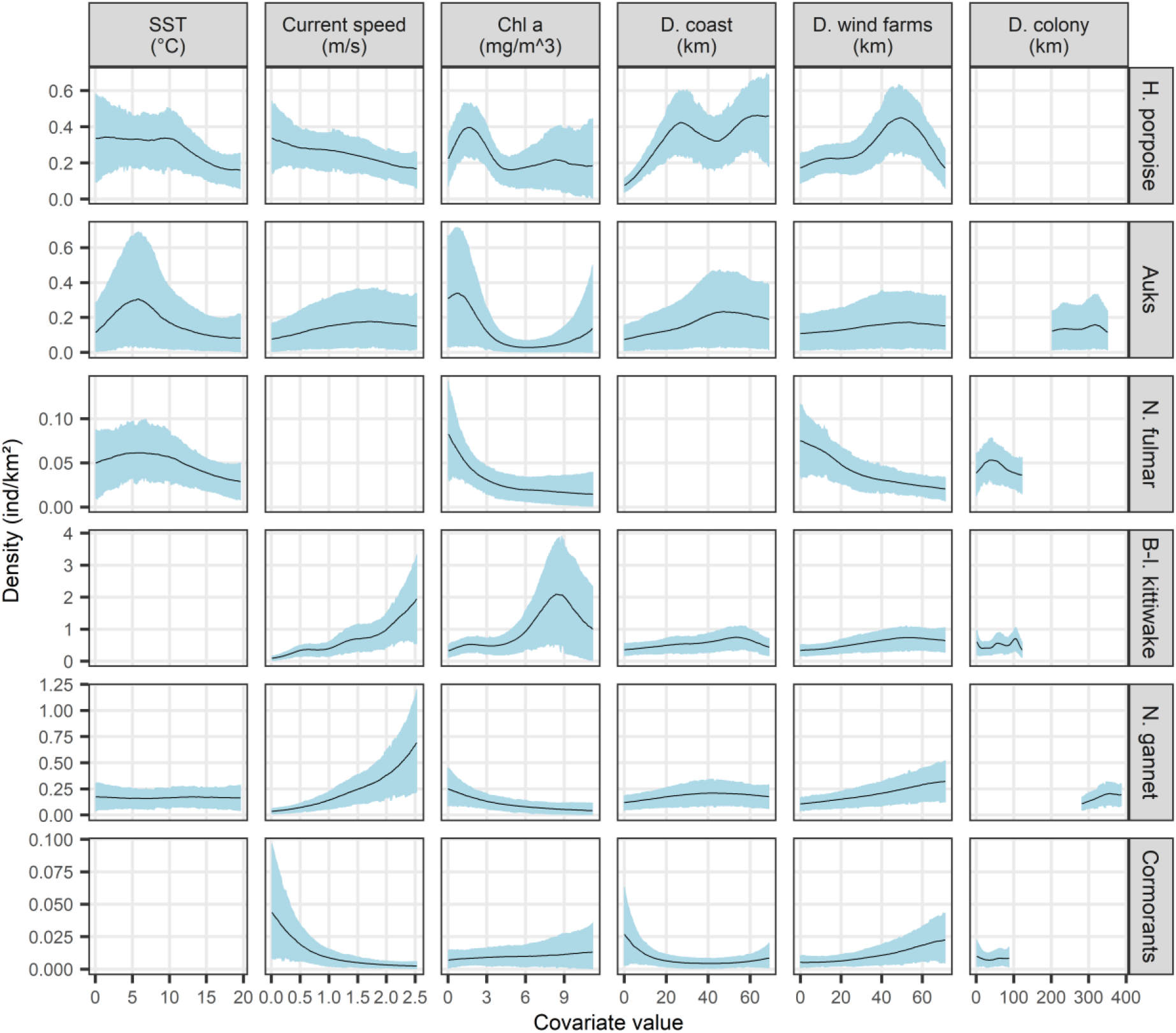
The average functional relationships between species and the selected variables. Solid lines represent the estimated smooth functions averaged over the entire period and the blue shaded regions show the approximate 95% confidence intervals (relationships obtained for each month are showed in Appendix G). The density of individuals is shown on the y-axis, where a zero indicates no effect of the covariate. A blank panel indicates that the variable was not selected in the model for this species group. (SST: sea surface temperature; Chl a: chlorophyll a concentration; D. coast: distance to coast; D. wind farms: distance to operational wind farms; D. colony: distance to the colony; B-l.: black-legged; H: harbour; N.: northern.)

As a result, harbour porpoises were not predicted in coastal areas but in offshore waters, with higher densities estimated further from the coasts (> 20 km, Figure 3A). Distribution patterns were relatively constant throughout the year but not the estimated densities and abundances. Estimated densities were lowest in summer (maximum density ≈ 0.1 individuals/km^2^) and highest in spring between March and May (maximum density ≈ 3-5 individuals/km^2^; Appendix H), with 2,500 individuals (80% Credible Interval: CI [970 - 4,200]) estimated in summer (August) and up to 25,000 individuals (CI [5,700 - 52,200]) estimated in spring (May, Appendix I).

**Figure 3.**
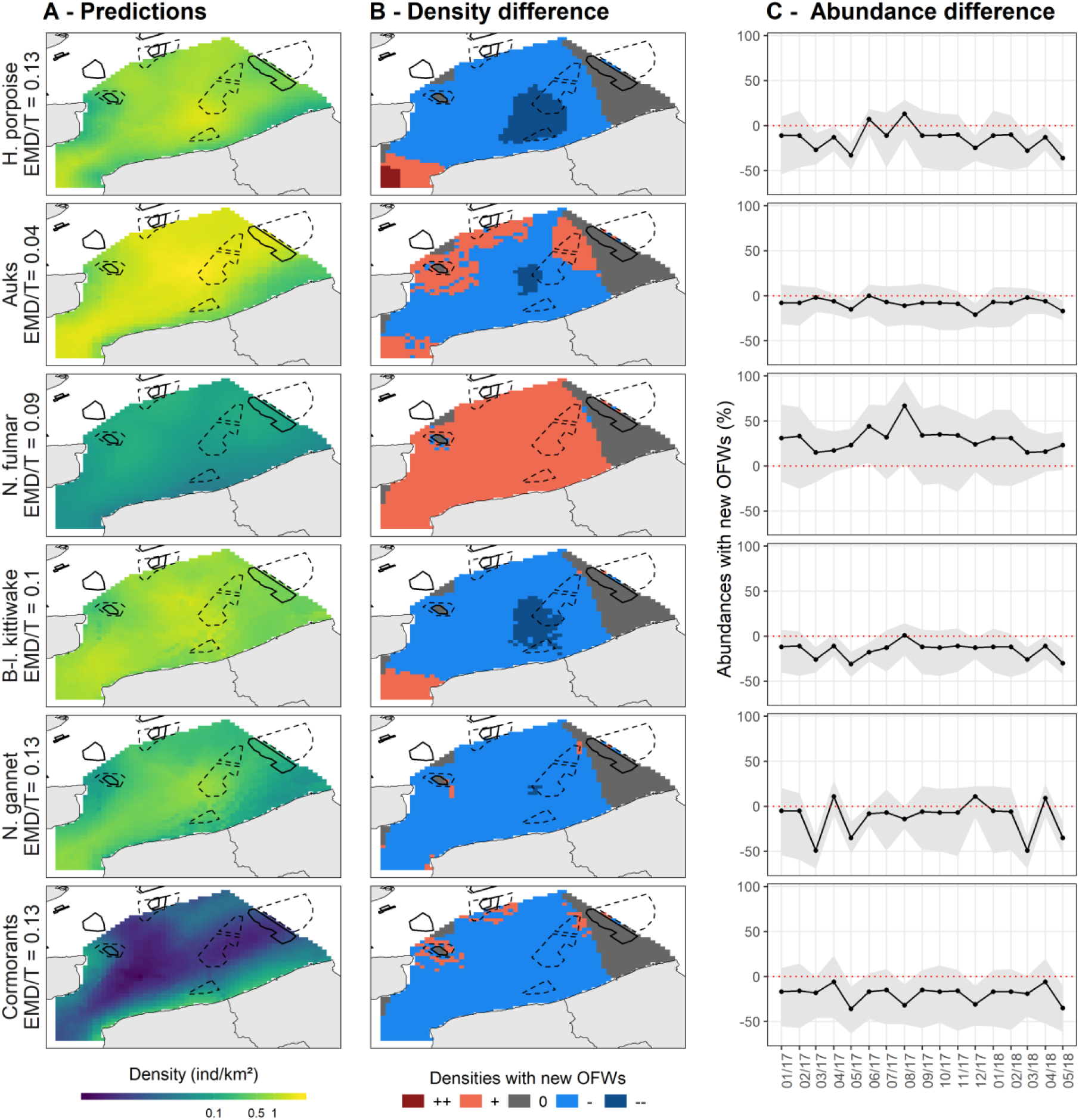
**(A) The average predicted densities obtained from the models in the southern North Sea.** Densities are averaged over the entire period. Monthly densities are shown in Appendix H. Black solid boxes represent the wind farms in operation and black dashed boxes the wind farms in development. **(B) The differences between counterfactuals and predicted densities.** Reds and oranges indicate that densities increased when the new OFWs were added; greys indicate that densities were not changed and blues indicate that densities decreased when the new OFWs were added. **(C) The monthly differences between counterfactual and predicted abundance estimates (number of individuals)**. The differences are given from January 2017 to May 2018. Positive values indicate that abundances increased when the new OFWs were added while negative values indicate that abundances decreased. Grey shaded regions represent confidence intervals.

##### Auks

The highest estimated densities of auks were associated with low chlorophyll *a* concentrations, large distances to the coast and colonies, low sea surface temperatures and average current speeds (Figure 2). Auks were predicted in offshore areas except in the north-west of the study area (near the English coast) and near the Belgian coast where they were almost absent (Figure 3A). They were most abundant between October and May (density > 1 individual/km2) and almost absent in summer (density < 0.1 individuals/km2, Appendix H) with about 100,000 individuals estimated in March (CI [14,000 - 260,000]) and only 100 individuals estimated in August (CI [8 - 250]; Appendix I).

##### Northern fulmars

The highest estimated densities of fulmars were associated with low chlorophyll *a* concentrations, low sea surface temperatures and short distances to colonies (Figure 2). Fulmars were distributed with high densities following a south-west / north-east axis that passed through the Dover Strait, especially in spring (March-April; Appendix H). Overall, abundances were quite low, reaching a maximum of 3,100 individuals in March (CI [150 – 8,850]) and a minimum of 300 (CI [10 - 570]; Appendix I) in August.

#### 3.2.2. Species abundant in winter

##### Black-legged kittiwakes

The highest estimated densities of black-legged kittiwakes were associated with high chlorophyll *a* concentrations, high current speeds and large distances to the coast (Figure 2). Kittiwakes were predicted to occur in the whole study area (Figure 3A) with an east-west shift in their distribution pattern over the year. The lowest abundance was estimated in August (≈ 4,700 individuals, CI [380 - 8,800]) and the highest in November (≈ 23,000 individuals; CI [750 - 65,600]; Appendix I).

#### Species with generally stable abundances all year round

##### Northern gannets

The highest estimated densities of northern gannets were associated with high current speeds, low chlorophyll *a* concentrations and large distances to the coast and colonies (Figure 2). Gannets were mostly predicted to occur in the centre of the study area (Figure 3A). They were more abundant in December and March with up to 10,000 individuals (CI [950 – 35,000]) and down to 3,000 - 4,000 individuals during the other months (Appendix I).

##### Cormorants

The highest estimated densities of cormorants were close to the colonies in shallow waters associated with low current speeds (Figure 2). Their resulting distribution was very coastal, predicted to be mainly along the French and Belgian coasts (Figure 3A) with little variation throughout the year. They were most abundant in March (≈ 2,700 individuals, CI [10 – 9,000], Appendix I).

### 3.3. Prospective effect of wind farm activity

The relationships between predicted densities and distance to operational OFWs varied between species groups (Figure 2). Predicted densities of northern fulmars were higher at short distances to OFWs while densities of auks were insensitive to distance to OFWs. In contrast, the highest densities of kittiwakes, cormorants, porpoises and gannets were found at distances larger than 40 km from OFWs.

The estimated distribution of all species in the eastern and north-western part of the study area remain unchanged when predictions from the prospective scenario, wherein all planned OFWs were assumed to be operating, are taken into account (Figure 3B). For porpoises, kittiwakes, gannets and cormorants, densities would be lower, on average, in the centre of the study area, particularly near the planned OFWs. Densities of fulmars would however be higher in this central area. The predicted change in auk distribution was more contrasted: densities were predicted to be lower in the centre of the study area but higher in the northern and western parts. The EMD values were relatively stable for auks, indicating little variation in the monthly distributions, whereas for all other species the EMD variations were larger, indicating larger monthly variations in distributions (Figure 4).

**Figure 4.**
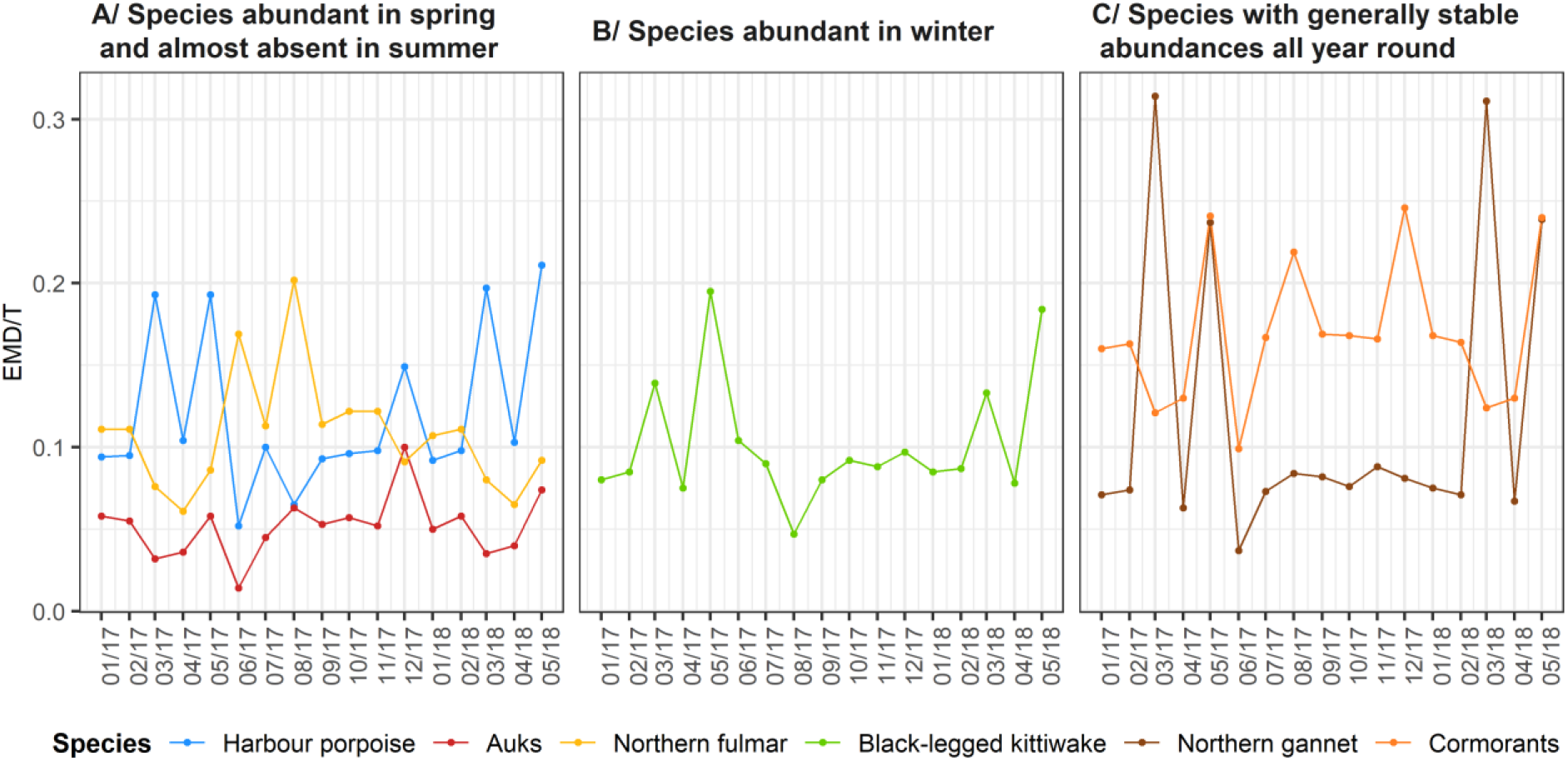
The earth mover’s distance thresholds (EMD/T) per species per month. The EMD is calculated as the distance between the model and counterfactual prediction. The EMD values are then standardised by dividing them by the threshold distance (here 20 km) resulting in the EMD/T values (see R-package “move”; Kranstauber et al., 2018). A value of 0 indicates a perfect match between the model and counterfactual predictions.

If the planned OFWs were operating, the abundance of fulmars would increase significantly (between +20% and +70%; Figure 3C), all else being equal, while the abundance of all other species would decrease (down to −50% for gannets), although the decrease would be less for auks (maximum −20%). There are two months during the time period where abundances of porpoises and gannets slightly increase (+10%; Figure 3C).

## 4. Discussion

For the year 2017/2018, using an approximate BAG method, we highlighted high intra-annual variability in distribution patterns with species mostly present in spring, or in winter or all year round. To minimise the impacts of OFWs currently planned in the area, construction should consider that species are not equally distributed within a year, and consequently take place in areas and seasons during which animals are present in the lowest densities. Our results highlight summer as the season where seabirds and harbour porpoises are present in the lowest densities in the area. We also highlight a potential change in species distribution after the planned OFWs come into operation, suggesting potential long-term effects on marine megafauna that need to be considered in a marine spatial planning context.

### 4.1. Avoiding and minimising OFW impacts

The construction phase of an OFW poses the greatest threat to marine mammals because the noise generated by pile driving drives away the animals (Vallejo et al., 2018) and can induce hearing loss (Tougaard et al., 2005; Thomsen et al., 2006; Dähne et al., 2013). During the operational phase, the noise levels generated by the turbines and maintenance activities are lower than those during pile driving and are unlikely to affect marine mammals (Madsen et al., 2006). The operational phase is the most hazardous for seabirds because of fatal collisions with wind turbines, particularly for gannets (Furness et al. 2013).

In order to meet the mitigation objectives of avoidance and minimisation (Kiesecker et al., 2010), we recommend that wind farms be established in areas of low seabird densities to minimise the risk of collisions during the operational phase, and that the construction phase be planned during periods of low porpoise densities. Harbour porpoises have an offshore distribution and were absent along the coast line. They appeared to prefer cool waters, which may explain their decreased abundance in summer compared to winter and spring. It is likely that they migrate to the central and northern parts of the North Sea in summer (Benke et al., 2014; McClellan et al., 2014). Most seabirds were predicted to be more abundant further offshore with monthly variations in their abundances with the exception of cormorants, which had a coastal distribution. This is because non-breeding cormorants feed within a radius of 35 km around their colonies and resting sites (Gremillet, 1997). Auks and fulmars were mostly abundant in spring and winter and absent from the study area in summer. This is because auks start migrating to their wintering grounds between May and October (Heath et al., 2000; Fort et al., 2013) and fulmars between June and November (Cadiou et al., 2004; Lambert et al., 2017). Kittiwakes were abundant in winter, which is when they are concentrated around breeding colonies, and less abundant in summer when they dispersed far from colonies (Wernham et al., 2002). The highest abundances of northern gannets were estimated at the beginning of spring, the prenuptial period during which many individuals cross the Dover Strait (Fort et al., 2012). Individuals observed between April and September were probably immature birds because mature individuals breed outside of the study area (Fort et al., 2012; Kubetzki et al., 2009; Wakefield et al., 2013).

We are aware that we sampled only a few months of the year and we cannot represent fixed monthly or seasonal distribution patterns for the studied species over multiple years. However, our model, which included month-varying relationships (modelled as cyclical) and time-varying spatial effects (modelled as random walk; see supplementary material) allowed to predict species distributions for each month, thereby illustrating intra-annual variation. Using this model, we would recommend (i) that the various OFWs planned for development be constructed between mid-summer and early fall to minimise the impact on harbour porpoises, and (ii) that offshore areas in the centre of the study area be avoided to minimise the impact on seabirds. Given these recommendations, it appears that the two wind farms planned off the Belgian coast (between 2.5 and 2.8° E and 51.25 and 51.7° N; Figure 1) may be very harmful for seabirds - particularly auks, kittiwakes and gannets - during the operational phase. The construction phase of a wind farm takes between 1.5 and 2 years, depending, in part, on the number of wind turbines constructed (Cartensen et al., 2006). Some stages of the construction phase are noisier than others and last between 3 and 5 months (Cartensen et al., 2006; Brandt et al., 2011). Therefore, the time window proposed in the study would be compatible with the most noise-generating construction stages of the wind farms. In addition, weather conditions in the area are generally most favourable between mid-summer and fall, which could facilitate the construction phase and limit its duration.

Our recommendations for minimisation measures critically hinge on knowledge of seasonal abundance patterns for porpoises and of spatial distribution patterns for seabirds. Carrying out regular surveys every month is very costly, and impractical over large areas. Here, we capitalised on a series of 6 surveys carried out between 2017 and 2018 to collect data on marine megafauna. These data were then used to calibrate a model that took into account spatio-temporal variations explicitly. In order to study mobile species, monitoring over a broad area is required which in turn limit temporal replicates. With 6 surveys spread over a whole year, we highlighted intra-annual variations in the species distribution allowing to identify the period and areas where the impact would be the lowest while with an average annual distribution, we would not have been able to do it. This study therefore highlights how critical spatial and seasonal variations in abundance are to marine spatial planning.

### 4.2. Prospective analysis of the impact of OWFs on marine megafauna

In this study, we have tried to apply the approach proposed by Methratta (2020), consisting in recording species sightings at different distances from wind farms and to include the distance to the wind farms in the models. We thus estimated relationships between the sightings and the distance to the nearest OFW as well as with variables that characterise the environment. The results are consistent with the known ecology of the species. The transect design carried out during the surveys does not strictly follow the BAG design because it is not a sampling at increasing distances from the wind farms but rather a sampling that allows the area to be optimally gridded with an aircraft (limitation of transit time with optimal coverage). We have therefore adapted the approach proposed by Methratta (2020) to the aerial survey methodology. We highlighted the effect of the OFW present in the area and use the obtained relationships to predict the distribution that species would have if these relationships were maintained with the OFWs currently planned in the area.

The estimated relationships between animal distributions and the planned OFWs must be interpreted with caution: they are not causal, nor do they arise from direct observations of behavioural responses to OFWs. Instead they could reflect the effect of other variables (*e.g*. environmental) not considered in the models. The models predicted the highest densities of fulmars at short distances to operational OFWs, whereas for all other groups save auks the highest densities were at large distances. Predicted densities of auks were unaffected by the distance to OFWs. Furness et al. (2013) showed that gannets are sensitive to the presence of OFWs and would avoid them. This is supported in our results by the predictions of lower gannet densities near the operating OFWs in the area. Likewise, Vallejo et al. (2018) showed a constant abundance of common guillemot (an auk species) regardless of the development phases of an OFW. Garthe and Hüppop (2004) have argued that cormorants, kittiwakes and fulmars were among the least sensitive species to the presence of OFWs. Porpoises seem to avoid an OFW area during the construction phase but can become as abundant during the operational phase as before construction started (Madsen et al., 2006; Brandt et al., 2011; Vallejo et al., 2018). Here, densities of harbour porpoises were lower in the east, near the Belgian wind farm, than in the centre of the study area, this may be due to the construction of an extension around the Belgian wind farm during the survey. Our model seems to be able to predict an avoidance of the construction area as demonstrated by Vallejo et al. (2018).

The prediction of seabird and harbour porpoise distributions based on the hypothetical scenario in which all planned OFWs were fully operational highlighted changes in species distribution. These predictions can be compared to future monitoring data, putting the model’s predictive ability to a true test and ultimately facilitating the improvement of predictive models. Northern fulmar abundance was predicted to increase near the new OFWs while the abundance of the other groups decreased, which suggests an avoidance of OFWs. These predicted changes in abundance and distribution clearly suggest that new OFWs in the study area would impact marine megafauna when construction is finalised and they become operational. To minimise this impact, we recommend that the number of OFWs constructed in the coming years be limited, and that the new OFWs be spaced more widely apart, avoiding high concentrations of farms in a small area. The construction of OWFs in various areas should also be staggered temporally so that the disruption occurs only in a limited area at a time, allowing animals, especially marine mammals, areas of refuge in which they can continue foraging, breeding, etc. undisturbed.

Our predictions are borne out of the best available evidence for the study area, and rely on a state-of-the-art modelling approach that takes into account seasonal and spatial variations. Regular monitoring of the study area in the future can ground-truth these predictions. Our prospective approach allows for the cumulative effect of the commissioning of multiple OFWs (either operating or planned) on marine megafauna distribution to be quantified, in contrast to the DEPONS model (Nabe-Nielsen et al., 2018), which assesses the cumulative impact of noise on harbour porpoise distribution during the construction phase of multiple OFWs. The approach and results we put forward in this study suggest additional impacts of OWFs than those determined using the DEPONS model. The DEPONS model considers the cumulative impact of noise from OFW projects built simultaneously or sequentially, while our approach targets various species, not only those impacted by noise, and assesses the long-term impacts on species distributions once planned OFWs are operational, and not only the impacts during the construction phase. These two approaches complement one another and are both crucial to mitigation efforts and marine spatial planning objectives with regards to the impacts of OFWs at the population scale since they individually concern the construction and operational phases.

Marine species are subject to anthropogenic pressures beyond those resulting from OFWs, and in the North Sea those pressures are many. The southern North Sea is crossed by a shipping lane frequently used by more than 400 commercial ships (1/4 of the world traffic), many ferries as well fishing and recreational vessels, which certainly have an impact on marine megafauna (Halpern et al., 2008). For example, Wisniewska and colleagues (2018) have shown that porpoises make deeper dives and interrupt their foraging and echolocation activities due to the noise of boats. Fishing also affects seabird and marine mammal populations through the alteration of food supplies and bycatch incidents in fishing gear (Tasker et al., 2000; Read et al., 2006). The cumulative impacts of boat noise, fishing and wind farm activities could have negative long-term consequences on marine megafauna. These impacts are important to consider in the context of marine spatial planning in order to establish and maintain an organised maritime area that takes the interactions between users (both professional and private) into account and balances the need for maritime development and activities with the protection of the environment (Douvere, 2008; European Parliament and Council Directive 2014/89/EU). The better we understand the impacts of anthropogenic activities – individually and cumulatively, on a specific and broad scale – the more effectively we can reconcile development and environmental conservation and make valuable suggestions for marine spatial planning.

## 5. Conclusion

The southern North Sea is subjected to intense human activities that include high maritime traffic (commercial, fishing and tourist vessels) as well as a high concentration of OFWs. The region is also densely populated by seabirds and harbour porpoises. It is crucial that we understand how marine megafauna use their environment and how they respond to maritime activities in order to 1) anticipate the effects of anthropogenic pressures, both current and future, on the animals that inhabit this region and 2) to make a guided effort to mitigate them. With mobile species, it is important to define how their distribution and abundance may change over time. Our study illustrates how the regular monitoring of marine megafauna in a relatively small area over several seasons enables the prediction of monthly distributions and abundances and changes thereof under hypothetical scenarios of increased human activities. Our study also illustrates how the BAG design can be used on aerial survey data. In addition to being promising for fisheries surveys, the BAG design approach has shown promising results in assessing the impact of wind farms on large predators such as cetaceans and seabirds.

One of the objectives of the European Union Member States is to contribute to the sustainable development of energy sectors at sea, maritime transport, and the fisheries and aquaculture sectors, but their role also includes ensuring the preservation and protection of the environment. This study aims to inform marine spatial planning decisions regarding these developments so that they have the least possible impact on marine species. Our goal was to reconcile the development of wind farm activity with the conservation of marine megafauna species by anticipating the potential impacts the existing and planned developments would have on various species using the best available evidence. While a prospective assessment helps in planning and decision making, it remains paramount to ensure that regular and accurate monitoring of the study area continues in order to ground-truth predictions and revise models accordingly both for future marine spatial planning and mitigation purposes.

## Supporting information

Supplementary material of the manuscript

## Data accessibility

Data are available via the OBIS SEAMAP website (http://seamap.env.duke.edu/).

## Acknowledgments

We thank above all the *Direction Générale de l’Energie et du Climat*, the *Agence Française pour la Biodiversité* and the CEREMA that enabled us to carry out this study. Surveys would not have been possible without the investment and flexibility of the Pixair Survey team that conducted the flight sessions: Jean-Jérôme Houdaille, Pierre Chevé and Christian Morales; and the team of observers: Cécile Dars (Pelagis), Ghislain Dorémus (Pelagis), Marc Duvilla (LPO Normandie), Sophie Laran (Pelagis), Morgane Perri (Al Lark) and Olivier Van Canneyt (Pelagis). We would also like to thank the air traffic control agencies that facilitated the access to the study area: the DGAC and the Air Navigation Service (SNA NORD) for France; BELGOCONTROL and the Koksijde Control Tower for Belgium; the Luchtverkeersleiding Nederland for the Netherlands; and NATS for England. We thank Carin Reisinger for proofreading the manuscript and contributing to its improvement.

## Appendices

Appendix A. Effort carried out during each flight session.

Appendix B. Comparison of harbour porpoise abundance estimations using conventional distance sampling and density surface modelling methods.

Appendix C. Colonies of seabirds located near the study area.

Appendix D. Details about the model used in the study.

Appendix E. Average monthly distributions of environmental variables used in the density surface modelling.

Appendix F. Sightings recorded during each flight session.

Appendix G. The monthly functional relationships between species and the selected variables.

Appendix H. Monthly predicted densities in the southern North Sea.

Appendix I. Monthly abundances in number of individuals estimated from the model (A) and the counterfactuals (B).

## References

Benke, H., Bräger, S., Dähne, M., Gallus, A., Hansen, S., Honnef, C.G., … & Narberhaus, I. (2014). Baltic Sea harbour porpoise populations: status and conservation needs derived from recent survey results. Marine Ecology Progress Series, 495: 275–290. doi: https://doi.org/10.3354/meps10538.

Bergström, L., Sundqvist, F., and Bergström, U. (2013). Effects of an offshore wind farm on temporal and spatial patterns in the demersal fish community. Marine Ecology Progress Series, 485: 199–210. doi: https://doi.org/10.3354/meps10344.

Bergström, L., Kautsky, L., Malm, T., Rosenberg, R., Wahlberg, M., Capetillo, N.Å., & Wilhelmsson, D. (2014). Effects of offshore wind farms on marine wildlife - a generalized impact assessment. Environmental Research Letters, 9(3): 034012. doi:10.1088/1748-9326/9/3/034012.

Board, O.S., & National Research Council. (2005). Marine mammal populations and ocean noise: determining when noise causes biologically significant effects. National Academies Press.

Brandt, M.J., Diederichs, A., Betke, K., & Nehls, G. (2011). Responses of harbour porpoises to pile driving at the Horns Rev II offshore wind farm in the Danish North Sea. Marine Ecology Progress Series, 421: 205–216. doi: https://doi.org/10.3354/meps08888.

Buckland, S.T., Anderson, D.R., Burnham, H.P., Laake, J.L., Borchers, D.L. & Thomas, L. (2001). Introduction to distance sampling: Estimating abundance of biological populations. Oxford University Press, Oxford.

Cadiou, B., Pons, J-M., & Yésou, P. (2004). Oiseaux marins nicheurs de France métropolitaine: 1960–2000. Biotope Editions, Mèze, Hérault.

Carstensen, J., Henriksen, O.D., & Teilmann, J. (2006). Impacts of offshore wind farm construction on harbour porpoises: acoustic monitoring of echolocation activity using porpoise detectors (T-PODs). Marine Ecology Progress Series 321 : 295–308. doi:10.3354/meps321295.

Dähne, M., Gilles, A., Lucke, K., Peschko, V., Adler, S., Krügel, K.,… & Siebert, U. (2013). Effects of pile-driving on harbour porpoises (*Phocoena phocoena*) at the first offshore wind farm in Germany. Environmental Research Letters, 8(2): 025002. doi:10.1088/1748-9326/8/2/025002.

Degraer, S., Brabant, R., Rumes, B. & Vigin, L. (eds). 2018. Environmental Impacts of Offshore Wind Farms in the Belgian Part of the North Sea: Assessing and Managing Effect Spheres of Influence. Brussels: Royal Belgian Institute of Natural Sciences, OD Natural Environment, Marine Ecology and Management, 136 p.

Dormann, C. F., Elith, J., Bacher, S., Buchmann, C., Carl, G., Carré, G., … & Münkemüller, T. (2013). Collinearity: a review of methods to deal with it and a simulation study evaluating their performance. Ecography, 36(1): 27–46. doi: https://doi.org/10.1111/j.1600-0587.2012.07348.x

Douvere, F. (2008). The importance of marine spatial planning in advancing ecosystem-based sea use management. Marine policy, 32(5): 762–771. doi: https://doi.org/10.1016/j.marpol.2008.03.021.

Drewitt, A.L., & Langston, R.H. (2006). Assessing the impacts of wind farms on birds. Ibis, 148: 29–42. doi: https://doi.org/10.1111/j.1474-919X.2006.00516.x.

ESRI, (2016). ArcGIS - A Complete Integrated System Environmental Systems Research Institute, Inc., Redlands, California. <http://esri.com/arcgis>.

European Parliament and Council Directive 2014/89/EU of 23 July 2014 establishing a framework for maritime spatial planning. Available from: https://eur-lex.europa.eu/legal-content/EN-FR/TXT/?uri=CELEX:32014L0089&from=FR.

Foley, M. M., Halpern, B. S., Micheli, F., Armsby, M. H., Caldwell, M. R., Crain, C. M., … & Carr, M. H. (2010). Guiding ecological principles for marine spatial planning. Marine Policy, 34(5): 955–966. doi: https://doi.org/10.1016/j.marpol.2010.02.001.

Fort, J., Pettex, E., Tremblay, Y., Lorentsen, S. H., Garthe, S., Votier, S., … & Bearhop, S. (2012). Meta-population evidence of oriented chain migration in northern gannets (*Morus bassanus*). Frontiers in Ecology and the Environment, 10(5): 237–242. doi: https://doi.org/10.1890/110194.

Fort, J., Steen, H., Strøm, H., Tremblay, Y., Grønningsæter, E., Pettex, E., … & Grémillet, D. (2013). Energetic consequences of contrasting winter migratory strategies in a sympatric Arctic seabird duet. Journal of Avian Biology, 44(3): 255–262. doi: https://doi.org/10.1111/j.1600-048X.2012.00128.x.

Furness, R. W., Wade, H. M., & Masden, E. A. (2013). Assessing vulnerability of marine bird populations to offshore wind farms. Journal of environmental management, 119: 56–66. doi: https://doi.org/10.1016/j.jenvman.2013.01.025.

Garthe, S., & Hüppop, O. (2004). Scaling possible adverse effects of marine wind farms on seabirds: developing and applying a vulnerability index. Journal of applied Ecology, 41(4): 724–734. doi: https://doi.org/10.1111/j.0021-8901.2004.00918.x.

Gilles, A., Adler, S., Kaschner, K., Scheidat, M., & Siebert, U. (2011). Modelling harbour porpoise seasonal density as a function of the German Bight environment: implications for management. Endangered species research, 14(2), 157–169. doi: https://doi.org/10.3354/esr00344

Gilles, A., Viquerat, S., Becker, E.A., Forney, K.A., Geelhoed, S.C.V., Haelters, J., Nabe-Nielsen, J., Scheidat, M., Siebert, U., Sveegaard, S., van Beest, F.M., van Bemmelen, R. & Aarts, G. (2016). Seasonal Habitat-Based Density Models for a Marine Top Predator, the Harbour Porpoise, in a Dynamic Environment. Ecosphere, 7(6): 1–22. doi:10.1002/ecs2.1367.

Green, R.H. (1979). Sampling Design and Statistical Methods for Environmental Biologists. John Wiley and Sons: New York, NY.

Grémillet, D. (1997). Catch per unit effort, foraging efficiency, and parental investment in breeding great cormorants (*Phalacrocorax carbo*). ICES Journal of Marine Science, 54(4): 635–644. doi: https://doi.org/10.1006/jmsc.1997.0250.

Guisan, A., & Thuiller, W. (2005). Predicting species distribution: offering more than simple habitat models. Ecology letters, 8(9): 993–1009. doi: https://doi.org/10.1111/j.1461-0248.2005.00792.x.

Halpern, B.S., Walbridge, S., Selkoe, K.A., Kappel, C.V., Micheli, F., D’agrosa, C., … & Fujita, R. (2008). A global map of human impact on marine ecosystems. Science, 319(5865): 948–952. doi: 10.1126/science.1149345.

Harwood, J. & King, S. (2014). Interim PCoD v1.1: a ‘how to’ guide, 28 p.

Hastie, T., & Tibshirani, R. (1986). Generalized Additive Models. Statistical Science, 3: 297–313.

Heath, M.F., Borggreve, C., & Peet, N. (2000). European bird populations : estimates and trends. BirdLife conservation series, 99-2048818-6: 10. BirdLife International, Cambridge.

Isaac, N.J., Jarzyna, M.A., Keil, P., Dambly, L.I., Boersch-Supan, P.H., Browning, E., … & Jarvis, S. (2019). Data Integration for Large-Scale Models of Species Distributions. Trends in ecology & evolution. doi: 10.1016/j.tree.2019.08.006.

Kiesecker, J. M., Copeland, H., Pocewicz, A., & McKenney, B. (2010). Development by design: blending landscape-level planning with the mitigation hierarchy. Frontiers in Ecology and the Environment, 8(5): 261–266. doi: https://doi.org/10.1890/090005.

Kranstauber, B., Smolla, M., & Safi, K. (2017). Similarity in spatial utilization distributions measured by the earth mover’s distance. Methods in Ecology and Evolution, 8(2): 155–160. doi: 10.1111/2041-210X.12649.

Kranstauber, B., Smolla, M., & Scharf, A.K. (2018). move: Visualizing and Analyzing Animal Track Data. R package version 3.1.0. https://CRAN.R-project.org/package=move.

Kubetzki, U., Garthe, S., Fifield, D., Mendel, B., & Furness, R. W. (2009). Individual migratory schedules and wintering areas of northern gannets. Marine Ecology Progress Series, 391: 257–265. doi: https://doi.org/10.3354/meps08254.

Lambert, C., Pettex, E., Dorémus, G., Laran, S., Stéphan, E., Van Canneyt, O. & Ridoux, V. (2017). How does ocean seasonality drive habitat preferences of highly mobile top predators? Part II: The eastern North-Atlantic. Deep Sea Research Part II: Topical Studies in Oceanography, 141: 133–154. doi: https://doi.org/10.1016/j.dsr2.2016.06.011.

Lambert, C., Authier, M., Doremus, G., Gilles, A., Hammond, P., Laran, S., … & Van Canneyt, O. (2019). The effect of a multi-target protocol on cetacean detection and abundance estimation in aerial surveys. Royal Society open science, 6(9): 190296. doi: https://doi.org/10.1098/rsos.190296

Legroux, N., Ponchon, A., Poirson, C. & Michel, S. (2017). Synthèse bibliographique sur les oiseaux migrateurs, nicheurs et hivernants dans le détroit du Pas-de-Calais, 1:173.

Madsen, P. T., Wahlberg, M., Tougaard, J., Lucke, K., & Tyack, P. (2006). Wind turbine underwater noise and marine mammals: implications of current knowledge and data needs. Marine Ecology Progress Series, 309: 279–295. doi:10.3354/meps309279.

McClellan, C.M., Brereton, T., Dell’Amico, F., Johns, D.G., Cucknell, A.C., Patrick, S.C., … & Votier, S.C. (2014). Understanding the distribution of marine megafauna in the English Channel region: identifying key habitats for conservation within the busiest seaway on earth. PloS one, 9(2): e89720. doi: https://doi.org/10.1371/journal.pone.0089720.

Mendel, B., Schwemmer, P., Peschko, V., Müller, S., Schwemmer, H., Mercker, M., & Garthe, S. (2019). Operational offshore wind farms and associated ship traffic cause profound changes in distribution patterns of Loons (*Gavia* spp.). Journal of environmental management, 231: 429–438. doi: https://doi.org/10.1016/j.jenvman.2018.10.053.

Methratta, E. T. (2020). Monitoring fisheries resources at offshore wind farms: BACI vs. BAG designs. ICES Journal of Marine Science, 77(3): 890–900. doi: https://doi.org/10.1093/icesjms/fsaa026.

Miller, D.L., Burt, M.L., Rexstad, E.A. & Thomas, L. (2013). Spatial Models for Distance Sampling Data: Recent Developments and Future Directions. Methods in Ecology and Evolution, 4: 1001–1010. doi:10.1111/2041-210X.12105.

Nabe-Nielsen, J., van Beest, F. M., Grimm, V., Sibly, R. M., Teilmann, J., & Thompson, P. M. (2018). Predicting the impacts of anthropogenic disturbances on marine populations. Conservation Letters, 11(5): e12563. doi: https://doi.org/10.1111/conl.12563.

Pitchford, J. L., Howard, V. A., Shelley, J. K., Serafin, B. J., Coleman, A. T., & Solangi, M. (2016). Predictive spatial modelling of seasonal bottlenose dolphin (Tursiops truncatus) distributions in the Mississippi Sound. Aquatic Conservation: Marine and Freshwater Ecosystems, 26(2), 289–306. doi: https://doi.org/10.1002/aqc.2547.

Read, A.J., Drinker, P., & Northridge, S.P. (2006). Bycatch of marine mammals in U.S. and global fisheries. Conservation Biology, 20: 163–169. doi: 10.1111/j.1523-1739.2006.00338.x

Roberts, J.J., Best, B.D., Dunn, D.C., Treml, E.A., & Halpin, P.N. (2010). Marine Geospatial Ecology Tools: An integrated framework for ecological geoprocessing with ArcGIS, Python, R, MATLAB, and C++. Environmental Modelling & Software, 25: 1197–1207. doi : https://doi.org/10.1016/j.envsoft.2010.03.029.

SAMMOA 1.0.4. Système d’Acquisition des données sur la Mégafaune Marine par Observations Aériennes, logiciel développé par l’UMS 3462 Pelagis et Code Lutin. http://www.observatoire-pelagis.cnrs.fr/publications/les-outils/article/logiciel-sammoa.

Santos, C.F., Agardy, T., Andrade, F., Crowder, L.B., Ehler, C.N., & Orbach, M.K. (2019). Major challenges in developing marine spatial planning. Marine Policy. doi: https://doi.org/10.1016/j.marpol.2018.08.032.

Strindberg, S., & Buckland, S. T. (2004). Zigzag survey designs in line transect sampling. Journal of Agricultural, Biological, and Environmental Statistics, 9(4): 443. doi: https://doi.org/10.1198/108571104X15601.

Tasker, M. L., Camphuysen, C. J., Cooper, J., Garthe, S., Montevecchi, W. A., & Blaber, S. J. (2000). The impacts of fishing on marine birds. ICES journal of Marine Science, 57(3): 531–547. doi: https://doi.org/10.1006/jmsc.2000.0714.

Thomas, L., Buckland, S.T., Rexstad, E.A., Laake, J.L., Strindberg, S., Hedley, S.L., … & Burnham, K.P. (2010). Distance software: design and analysis of distance sampling surveys for estimating population size. Journal of Applied Ecology, 47(1): 5–14. doi: https://doi.org/10.1111/j.1365-2664.2009.01737.x.

Thomsen, F., Lüdemann, K., Kafemann, R., & Piper, W. (2006). Effects of offshore wind farm noise on marine mammals and fish. Biola, Hamburg, Germany on behalf of COWRIE Ltd 62.

Tougaard, J. & Teilmann, J. (2005). Effects of the Horns Reef Wind Farm on harbour porpoises. Interim report to Elsam Engineering A/S for the harbour porpoise-monitoring program 2004, 23 p.

Vallejo, G.C., Grellier, K., Nelson, E.J., McGregor, R.M., Canning, S.J., Caryl, F.M., & McLean, N. (2017). Responses of two marine top predators to an offshore wind farm. Ecology and Evolution, 7(21): 8698–8708. doi: https://doi.org/10.1002/ece3.3389.

Vehtari, A., Gelman, A., & Gabry, J. (2017). Practical Bayesian model evaluation using leave-one-out cross-validation and WAIC. Statistics and computing 27(5): 1413–1432.

Wakefield, E.D., Bodey, T.W., Bearhop, S., Blackburn, J., Colhoun, K., Davies, R., … & Jessopp, M.J. (2013). Space partitioning without territoriality in gannets. Science, 341(6141): 68–70. doi: 10.1126/science.1236077.

Wernham, C. (2002). The migration atlas: movements of the birds of Britain and Ireland. T & AD Poyser.

Whitt, A. D., Dudzinski, K., & Laliberté, J. R. (2013). North Atlantic right whale distribution and seasonal occurrence in nearshore waters off New Jersey, USA, and implications for management. Endangered Species Research, 20(1), 59–69. doi: https://doi.org/10.3354/esr00486

Wikle, C., & Royle, J. (2002). Spatial statistical modeling in biology. Mississauga, ON: EOLSS Publishers Co. Ltd.

Wilber, D.H., Carey, D.A., & Griffin, M. (2018). Flatfish habitat use near North America’s first offshore wind farm. Journal of Sea Research, 139: 24–32. doi: https://doi.org/10.1016/j.seares.2018.06.004.

Wisniewska, D.M., Johnson, M., Teilmann, J., Siebert, U., Galatius, A., Dietz, R., & Madsen, P.T. (2018). High rates of vessel noise disrupt foraging in wild harbour porpoises (*Phocoena phocoena*). Proc. R. Soc. B, 285(1872): 20172314. doi: https://doi.org/10.1098/rspb.2017.2314.

Wood, S.N. (2006). On confidence intervals for generalized additive models based on penalized regression splines. Australian & New Zealand Journal of Statistics, 48: 445–464. doi: https://doi.org/10.1111/j.1467-842X.2006.00450.x.

