## Supplementary material of the manuscript for "Prospective modelling of operational offshore windfarms on the distribution of marine megafauna in the southern North Sea"

### Abstract

Intense development of Offshore Wind Farms (OFWs) has occurred in the North Sea with several more farms planned for the near future. These OFWs pose a threat to marine megafauna stressing the need to mitigate the impact of human activities. To help mitigating impacts, the Before After Gradient (BAG) design was proposed. We thus explored the use of the BAG method on megafauna sightings recorded at different distances from wind farms in the southern North Sea. We predicted intra-annual variability in species distribution, then correlated species distribution with the presence of operational OFWs and investigated the potential impact the operation of prospective OFWs may have on species distribution. Three patterns of intra-annual variability were predicted: species present most abundantly in spring, in winter or all year-round. We recommend that future OFW constructions be planned in summer and early fall to minimise impact on cetaceans and that offshore areas off northern France and Belgium be avoided to minimise impact on seabirds. Our prospective analysis predicted an increased or a decreased density with the operation of prospective OFWs. Prospective approaches, using e.g.

a BAG design, are paramount to inform species conservation as they can forecast the likely responses of megafauna to anthropogenic disturbances.

**Appendix A. Effort carried out during each flight session.** The black dotted lines delineate the study area and the blue solid lines represent the effort performed in each flight session. UK: "United Kingdom".

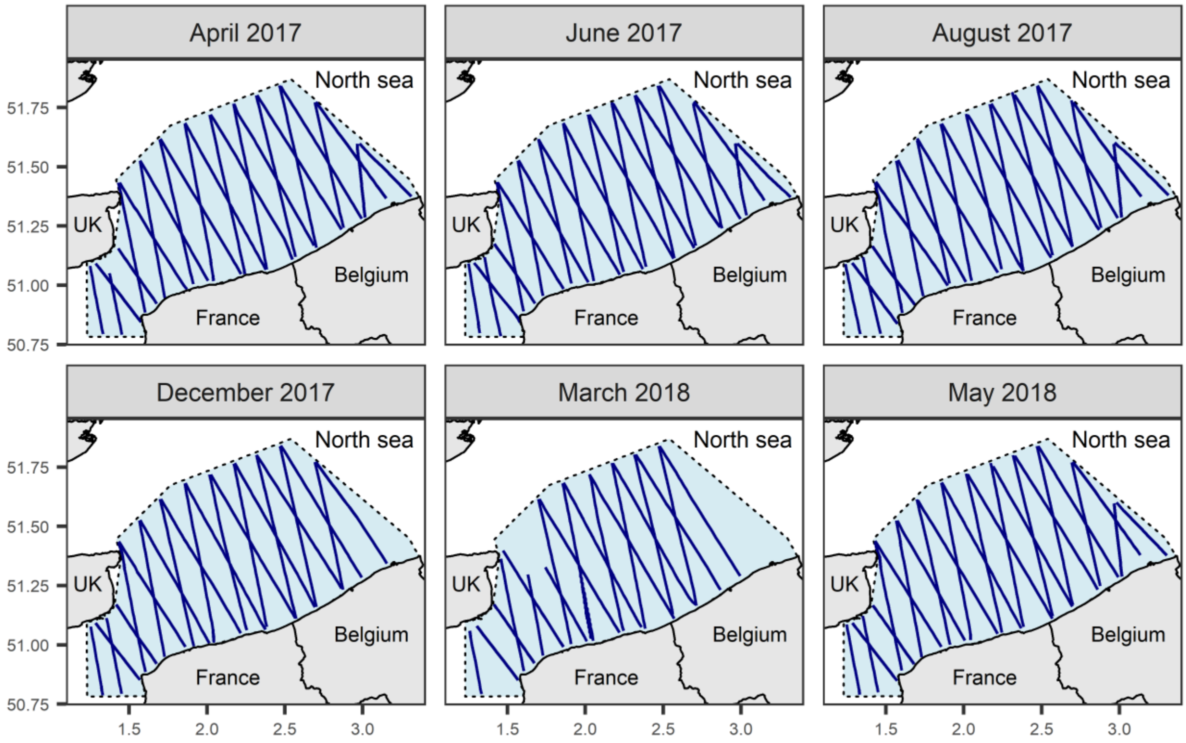

**Appendix B. Comparison of harbour porpoise abundance estimations using conventional distance sampling and density surface modelling methods.** Abundance estimates are shown for each month between January 2017 and May 2018. The grey shaded region represents confidence intervals. Confidence intervals are very tight around the abundance estimates when a flight session is associated with a certain month and are much wider when no flight session occurred during that month (the uncertainty is greater), particularly between September and November because the models were calibrated on the flight session in August and December. Red dots represent abundances and associated standard errors estimated from the conventional distance sampling method when a flight session was carried out.

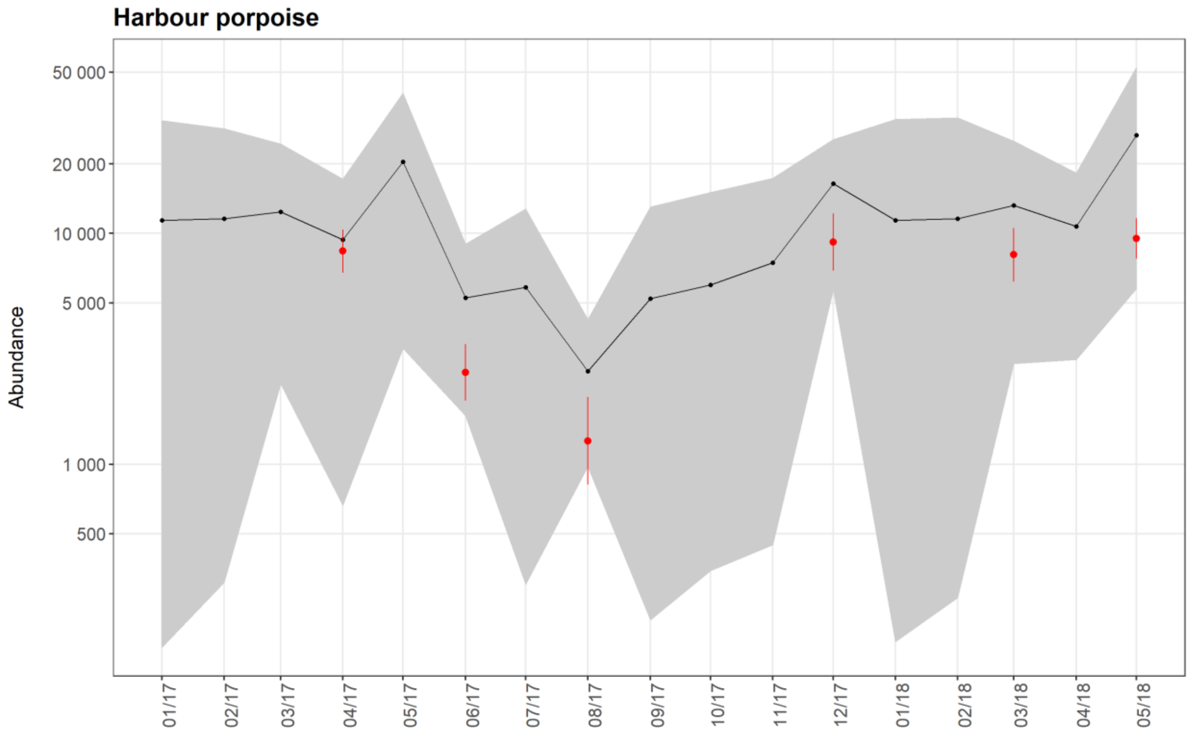

**Appendix C. Colonies of seabirds located near the study area.** These colonies were used to calculate the closest distance between the sightings and the colonies. The red dots represent the colonies; the dashed light blue zones represent the study area.

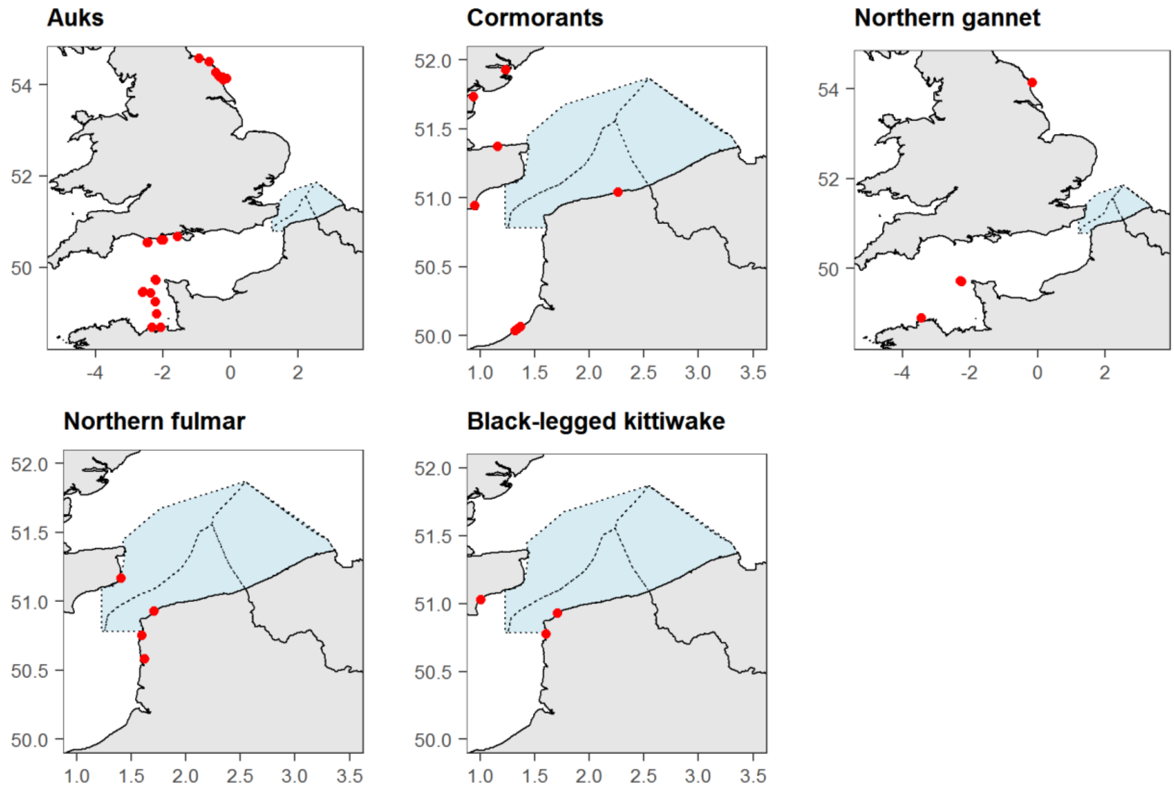

### Appendix D. Details about the model used in the study.

The model we used is a density surface model (Miller et al., 2013) to estimate the abundance of marine megafauna. Because the same area was repeatedly sampled over the course of a year, we adopted a spatio-temporal explicit approach to modelling. The model specification was:

$$\begin{cases} y_{imst} \sim \mathcal{NB}(\omega, \lambda_{imst}) \\ \log \lambda_{imst} = \log \text{Area}_{imt} + \alpha + \beta_m + \sum_{j=1}^{p} f_j^m X_j + \text{pixel}_{st} \\ \beta \sim \mathcal{GP}(\mathbf{0}, \sigma^2_{\text{season}} R) \end{cases}$$

where $i$ denotes the $i^{th}$ segment, $m$ the $m^{th}$ month of the year, $s$ the $s^{th}$ spatial pixel over the study area, and $t$ the $t^{th}$ survey session. Therefore, $y_{imst}$ corresponds to the number of animals detected on segment $i$ in month $m$ in pixel $s$ during session $t$. The parameter $\alpha$ represents the intercept. A binomial negative likelihood (with overdispersion parameter $\omega$) was assumed for the data as is customary with count data in ecology because of overdispersion. The $\beta$ parameters modelled monthly variations of the average density over the study area and this effect was assumed to be cyclical: for example, the average density in January was correlated with the average density in December and February but not with the average density in August. This information was incorporated into the R correlation matrix (the GP notation denotes a Gaussian process). An interaction with the month of the survey was included in the smoothing functions (cubic Beziers splines) so that relationships with the environment can vary by month. For each month $m$ and each predictor $j$, the environmental relationship between marine megafauna and predictors was modelled as:

$$\begin{cases} \forall\ (m, j),\ f_j^m(.) \sim \mathcal{GP}(\mu_j(.), \sigma_j^2 D) \\ \forall\ j,\ \mu_j(X_j) = \sum_{k=1}^{n_{\text{knots}}} \delta_k \text{BS}_k(x_{jk}) \end{cases}$$

where $\mu_j(.)$ is the mean relationship with predictor $j$ and is specified semi-parametrically with cubic Beziers splines with 20 equally spaced knots (Eilers & Marx, 2010). $D$ is a correlation matrix assuming a Matern correlation function of order 3/2 with the range parameter fixed to 1. Finally, a spatio-temporal effect ($pixel_{st}$) was included as a correlated random walk:

$$\forall t,\ \epsilon_t \sim \mathcal{GP}(\mathbf{0}, \sigma^2_{\text{spatial}} \Omega)$$

$$\forall s,\ \begin{cases} \text{pixel}_{st} = \epsilon_{st} & t = 1 \\ \text{pixel}_{st} = \rho \epsilon_{s(t-1)} + \epsilon_{st} \sqrt{1 - \rho^2} & t > 1 \end{cases}$$

Spatial dependence was incorporated into the $\Omega$ correlation matrix assuming a Matern correlation function of order 3/2 with the range parameter fixed so that two neighbouring pixels had a correlation of 0.5. This model is a hierarchical model that takes into account a time varying spatial effect, monthly density changes and possible monthly changes in environmental relationships over the study area. The parameter $\rho$ represents the temporal correlation (which takes into account the order of the flight sessions) while the matrix $\Omega$ represents the spatial correlation. This allows to take into account variations in density over time in the study area (contrary to the parameter $\beta$, time is decoupled from the month and its effect is not cyclical) and variations that would not be taken into account by the environmental covariates. The model was fitted using 'Stan v.2.17.3' software (Carpenter et al., 2017) called from the R software via the library 'RStan' (Stan model below; Stan Development Team, 2018). Weakly-informative normal priors were used (see Stan code below). All models took into account the sampled area associated with each segment as an offset. For seabirds, the sampled area was calculated as the segment length multiplied by twice the transect width (2 × 200 m), while for harbour porpoises, the sampled area was calculated as the segment length multiplied by twice the effective strip half-width (*esw*; Buckland et al., 2001). The *esw* is a distance estimated from the detection function, which depends on the perpendicular distance between the observation and the transect. It is determined so that the number of animals detected beyond this distance is equal to the number of undetected animals before this limit. Beforehand, 5% of the farthest sightings from the transect were discarded (Buckland et al., 2001). The detection function was then estimated by taking into account the observation conditions (sea-state, turbidity, flight session, subjective conditions, etc.) in order to correct the estimates according to these observation conditions ('Distance' R-package; Miller, 2017). Models using an increasing number of observation conditions were tested, the model with the lowest Akaike criterion (AIC; Anderson & Burnham, 2002) was selected and the *esw* was estimated. Detection of harbour porpoises was only affected by the flight session: an *esw* was estimated for each flight session accordingly (Figure S4.1).

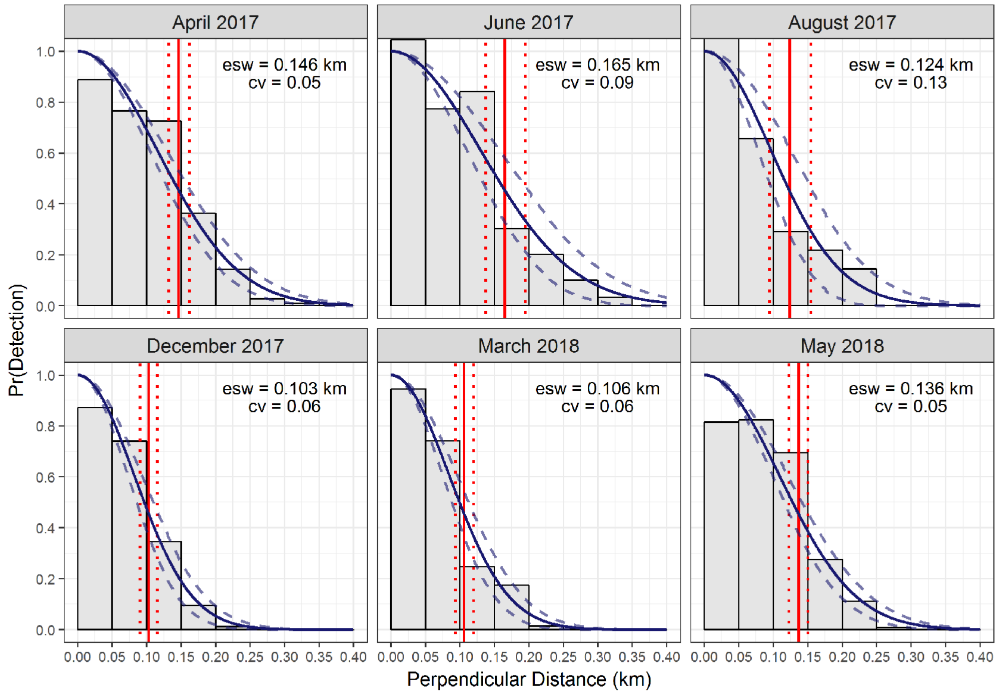


**Figure D.1. Detection functions estimated for each flight session for harbour porpoises.** Barplots represent the number of individuals observed depending on the perpendicular distance. Solid lines represent the estimated detection functions and dotted lines the confidence intervals.

A model selection procedure was used to select the best model for each species group. To avoid collinearity between variables, we first excluded combinations of variables with Spearman correlation coefficients greater than | 0.7 | (Dormann et al., 2013). The "Distance to wind farms in operation" for harbour porpoises and the "Distance to wind farms in operation" and "Distance to colony" for seabirds were always included in the fitted models to estimate the relationship between the number of individuals and the distance to the operational OFWs and to assess the effect of distance to the colonies on seabird distribution. For the other variables, the model selection procedure, based on the Widely Applicable Information Criterion (Vehtari et al., 2017), aimed to select the most predictive variables of species distribution by testing all combinations of variables. A maximum of five variables was selected for harbour porpoises and six variables for seabirds.

**Stan model code:**

```
data {
  int<lower = 1> n_cov;
```

```
  int<lower = 1> n_knot;
  int<lower = 1> n_cell;
  int<lower = 1> n_survey;
  int<lower = 1> n_obs[n_survey];
  int<lower = 0> cumul_obs[n_survey];
  int<lower = 1> n_obs_max;                        // for handling ragged arrays
  int<lower = 1, upper = 12> MONTH[n_survey];          // month of the survey
  int<lower = 1, upper = 7> SESSION[n_survey];         // time steps between surveys in pair of months
  int<lower = 0> Y[n_obs_max, n_survey];               // count data
  vector<lower = 0>[n_obs_max] EFFORT[n_survey];   // offset
  matrix[12, 12] chol_month;                       // Cholesky decomposition, lower triangular
  real prior_location_inter;
  real<lower = 0.0> prior_scale_inter;
  real<lower = 0.0> prior_scale_slope;
  vector<lower = 0.0>[4] prior_scale_sigma;
  vector<lower = 0.0>[n_survey] ESW_hat;
  vector<lower = 0.0>[n_survey] ESW_se;
  cov_matrix[n_survey] R;                              // correlation matrix for the esw
  matrix[n_cell, n_cell] chol_spatial;             // Cholesky decomposition, lower triangular
  matrix[n_obs_max, n_cell] W[n_survey];               // spatial weight matrix
  matrix[n_obs_max, n_cov] X[n_survey];                // design matrix
  matrix[n_obs_max, n_knot + 1] Z[n_survey, n_cov];  // Beziers-splines basis
  matrix[n_knot + 1, n_knot + 1] D;                    // Cholesky decomposition, lower triangular
  real<lower = 0.0> alpha;                             // concentration parameter of the Dirichlet distribution
}

parameters {
  real u_inter;
  vector[12] eps;
  vector<lower = 0>[4] tau;
  vector<lower = 0>[4] unscaled_sigma2;
  vector[n_survey] esw;
  vector[n_cell] u_spat[7];
  real<lower = -1.0, upper = 1.0> temporal_range;
  vector[n_knot + 1] unscaled_smooth[n_cov];
  vector[n_knot + 1] u_z[n_cov, 12];
  vector[n_cov] u_slope;
  real logit_omega;
  simplex[n_cov] psi[2];
}

transformed parameters {
  vector[n_survey] ESW;
  real intercept;
  vector[2] sigma_survey;
  vector[12] beta;
  vector[n_obs_max] linpred[n_survey];
```

```
  vector[n_cell] spatial[7];
  vector[n_cov] sigma_gp;
  vector[n_cov] slope;
  vector[n_cov] sigma_smooth;
  vector[n_knot + 1] z[n_cov, 12];
  vector[n_knot + 1] smooth[n_cov];
  real inv_omega;

  // overdispersion
  inv_omega = exp(1.5 * logit_omega);

  // effective strip width
  ESW = 2 * (ESW_hat + ESW_se .* esw);

  // overall intercept
  intercept = prior_location_inter + prior_scale_inter * u_inter;

  // slope
  slope = u_slope * prior_scale_slope;

  // survey-level variability
  sigma_survey[1] = prior_scale_sigma[1] * sqrt(unscaled_sigma2[1]/tau[1]);
  sigma_survey[2] = prior_scale_sigma[2] * sqrt(unscaled_sigma2[2]/tau[2]);

  // survey-specific: mean density
  beta = rep_vector(intercept, 12) + sigma_survey[1] * eps;

  // spatial effects: temporally correlated
  spatial[1] = u_spat[1]; // initial condition
  for(l in 2:7) {
    spatial[l] = temporal_range * spatial[l - 1] + sqrt(1 - square(temporal_range)) * u_spat[l];
  }

  // smooth functions
  for(j in 1:n_cov) {
    sigma_smooth[j] = prior_scale_sigma[3] * sqrt(psi[1, j] * unscaled_sigma2[3] / tau[3]);
    smooth[j] = sigma_smooth[j] * unscaled_smooth[j]; // overall mean of GP
    sigma_gp[j] = prior_scale_sigma[4] * sqrt(psi[2, j] * unscaled_sigma2[4] / tau[4]);
    for(l in 1:12) {
      z[j, l] = smooth[j] + sigma_gp[j] * u_z[j, l];
    }
  }

  // linear predictor
  for(k in 1:n_survey) {
    linpred[k] = rep_vector(log(ESW[k]) + beta[MONTH[k]] - 1.5 * logit_omega, n_obs_max) +
log(EFFORT[k]) + X[k] * slope + W[k] * (sigma_survey[2] * spatial[SESSION[k]]);
    for(j in 1:n_cov) {
```

```
      linpred[k] += Z[k, j] * z[j, MONTH[k]];
    }
  }
}

model {
  // over dispersion
  logit_omega ~ normal(0.0, 1.0);

  // scaled beta-2 prior for variances
  tau ~ gamma(1.0, 1.0);
  unscaled_sigma2 ~ gamma(0.5, 1.0);
  psi[1] ~ dirichlet(rep_vector(alpha, n_cov));
  psi[2] ~ dirichlet(rep_vector(alpha, n_cov));

  // Effective Stip Width
  esw ~ multi_normal(rep_vector(0.0, n_survey), R);

  // slope and intercept
  u_inter ~ normal(0.0, 1.0);
  u_slope ~ normal(0.0, 1.0);
  eps[1] ~ multi_normal_cholesky(rep_vector(0.0, 12), chol_month);

  // Spatial Effect
  for(l in 1:7) {
    u_spat[l] ~ multi_normal_cholesky(rep_vector(0.0, n_cell), chol_spatial);
  }

  // coefficients for spline basis
  for (j in 1:n_cov) {
    unscaled_smooth[j] ~ normal(0.0, 1.0);
    for (l in 1:12) {
      u_z[j, l] ~ multi_normal_cholesky(rep_vector(0.0, n_knot + 1), D);
    }
  }

  // Likelihood
  for (k in 1:n_survey) {
    for(i in 1:n_obs[k]) {
      target += neg_binomial_lpmf(Y[i, k] | inv_omega, exp(-linpred[k, i]));
    }
  }
}

generated quantities {
  real log_lik[sum(n_obs)];
  for(k in 1:n_survey) {
    for(i in 1:n_obs[k]) {
```

```
      log_lik[i + cumul_obs[k]] = neg_binomial_lpmf(Y[i, k] | inv_omega, exp(-linpred[k, i]));
    }
  }
}
```

### References


Anderson, D.R., & Burnham, K.P. (2002). Avoiding pitfalls when using information-theoretic methods. *The Journal of Wildlife Management,* 66(3): 912–918. doi: 10.2307/3803155.

Buckland, S.T., Anderson, D.R., Burnham, H.P., Laake, J.L., Borchers, D.L. & Thomas, L. (2001). *Introduction to distance sampling: Estimating abundance of biological populations*. Oxford University Press, Oxford.

Carpenter, B., Gelman, A., Hoffman, M.D., Lee, D., Goodrich, B., Betancourt, M.,… & Riddell, A. (2017). Stan: A Probabilistic Programming Language. *Journal of Statistical Software,* 76. doi:  10.18637/jss.v076.i01.

Dormann, C.F., Elith, J., Bacher, S., Buchmann, C., Carl, G., Carré, G.,... & Münkemüller, T. (2013). Collinearity: a review of methods to deal with it and a simulation study evaluating their performance. *Ecography,* 36(1): 27-46. doi: https://doi.org/10.1111/j.1600-0587.2012.07348.x.

Eilers, P.H.C. & Marx, B.D. (2010) Splines, Knots, and Penalties. WIREs Computational Statistics, 2010, 2, 637-653. doi: https://doi.org/10.1002/wics.125

Miller, D.; Burt, M.; Rexstad, E. & Thomas, L. (2013) Spatial Models for Distance Sampling Data: Recent Developments and Future Directions. Methods in Ecology and Evolution, 4, 1001-1010. doi: https://doi.org/10.1111/2041-210X.12105

Miller, D.L. (2017). Distance: Distance Sampling Detection Function and Abundance Estimation. R package version 0.9.7.  https://CRAN.R-project.org/package=Distance.

Stan Development Team. (2018). RStan: the R interface to Stan. R package version 2.17.3. http://mc-stan.org/.

Vehtari, A.; Gelman, A. & Gabry, J. (2017). Practical Bayesian Model Evaluation using Leave-One-Out Cross-Validation and WAIC. *Statistics and Computing*, 27: 1413-1432. doi: 10.1007/s11222-016-9696-4.

### Appendix E. Average monthly distributions of environmental variables used in the density surface modelling.

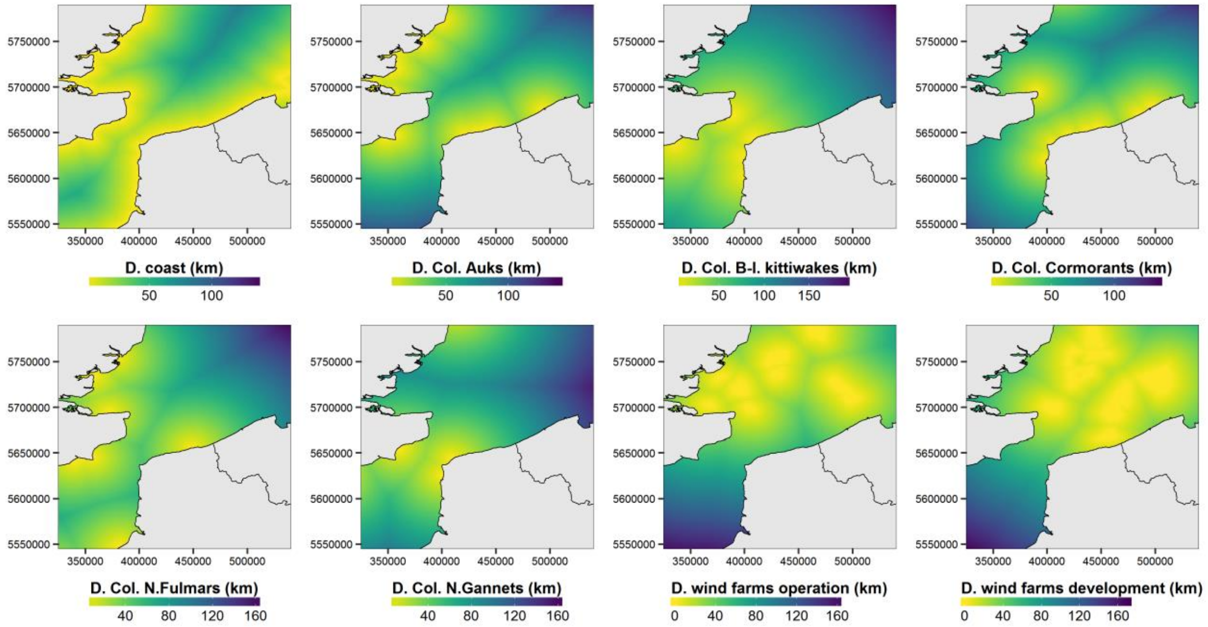


**Figure E.1. Distribution of static variables.** The distance to the coast was calculated in ArcGIS 10.3. The distance to the wind farms in operation was calculated in ArcGIS 10.3 from the wind farm perimeters provided by http://www.marineatlas.be/en/data for Belgium and by https://www.thecrownestate.co.uk/en-gb/resources/maps-and-gis-data/ for the United Kingdom. The distance to the wind farms in development was calculated in ArcGIS 10.3 from the wind farm perimeters provided by https://www.4coffshore.com/offshorewind/. The distance to the colonies was calculated in ArcGIS 10.3 from the location of the colonies (Appendix 2).

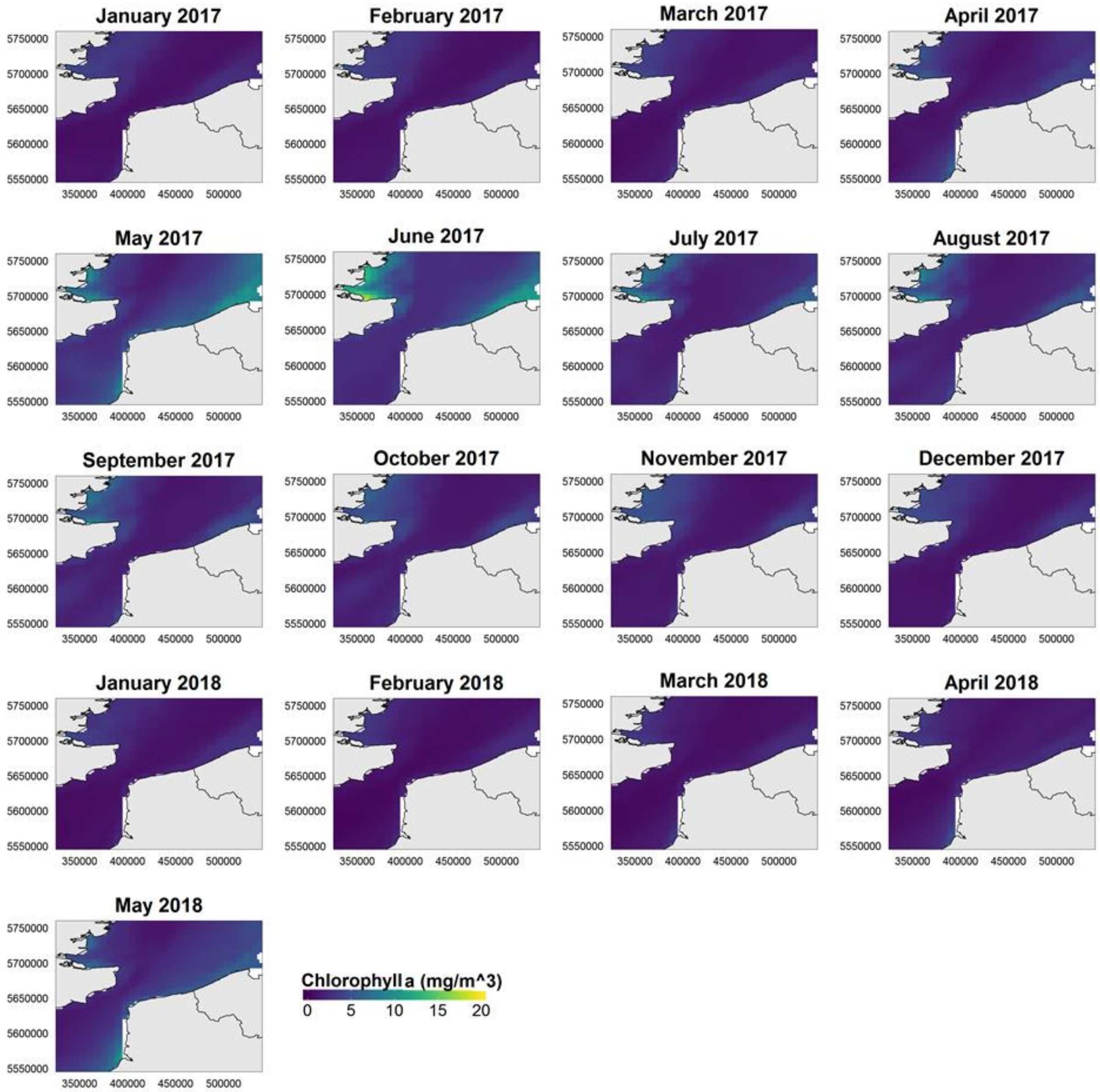


**Figure E.2. Monthly distribution of chlorophyll *a* concentration.** Average distribution for all months between January 2017 and May 2018. The concentration of surface chlorophyll was extracted from the ECOMARS 3D model from IFREMER (https://marc.ifremer.fr/resultats/).

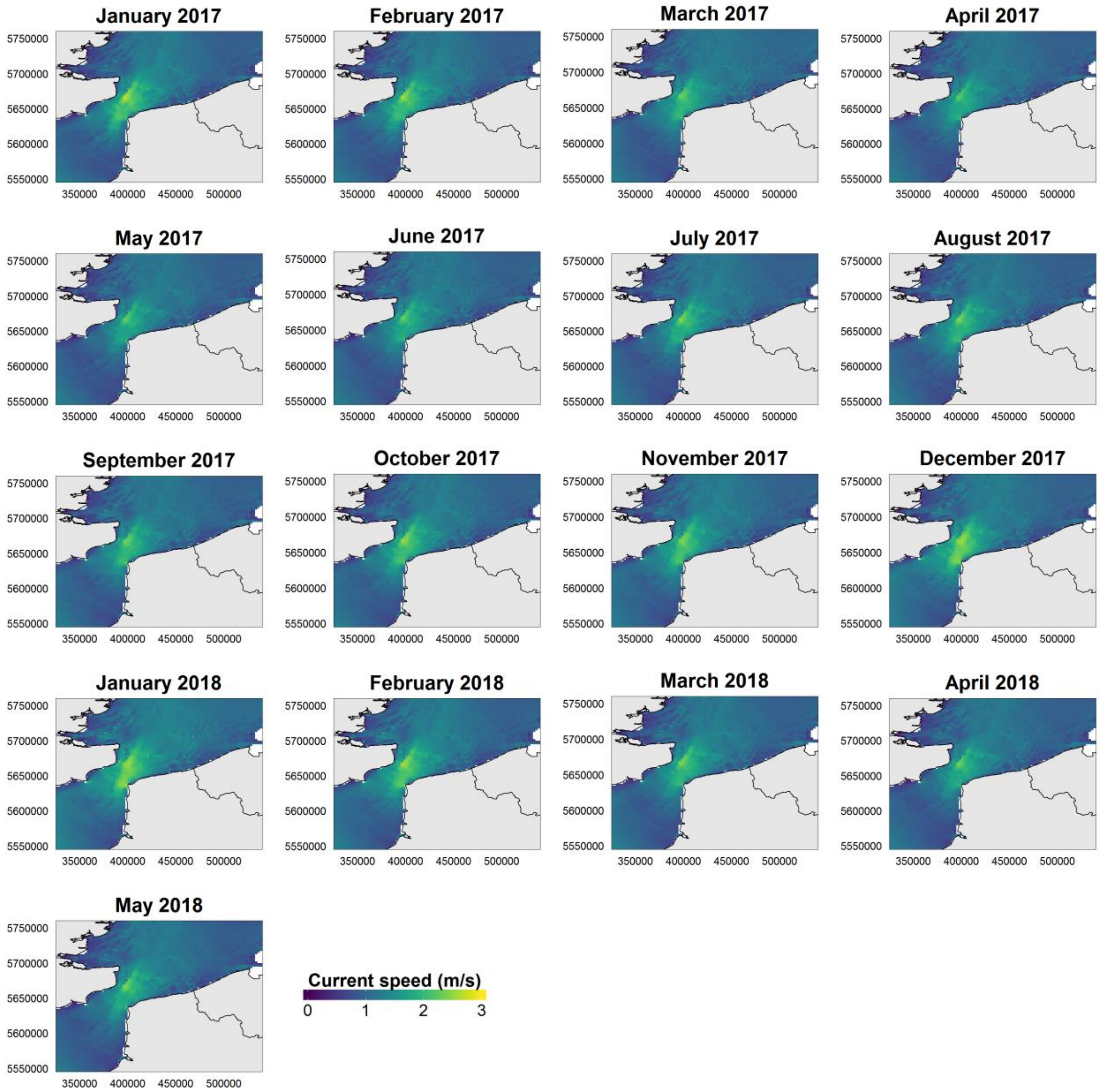


**Figure E.3. Monthly distribution of current speed.** Average distribution for all months between January 2017 and May 2018. The current was calculated from the MARS 2D model from IFREMER (https://marc.ifremer.fr/resultats/).

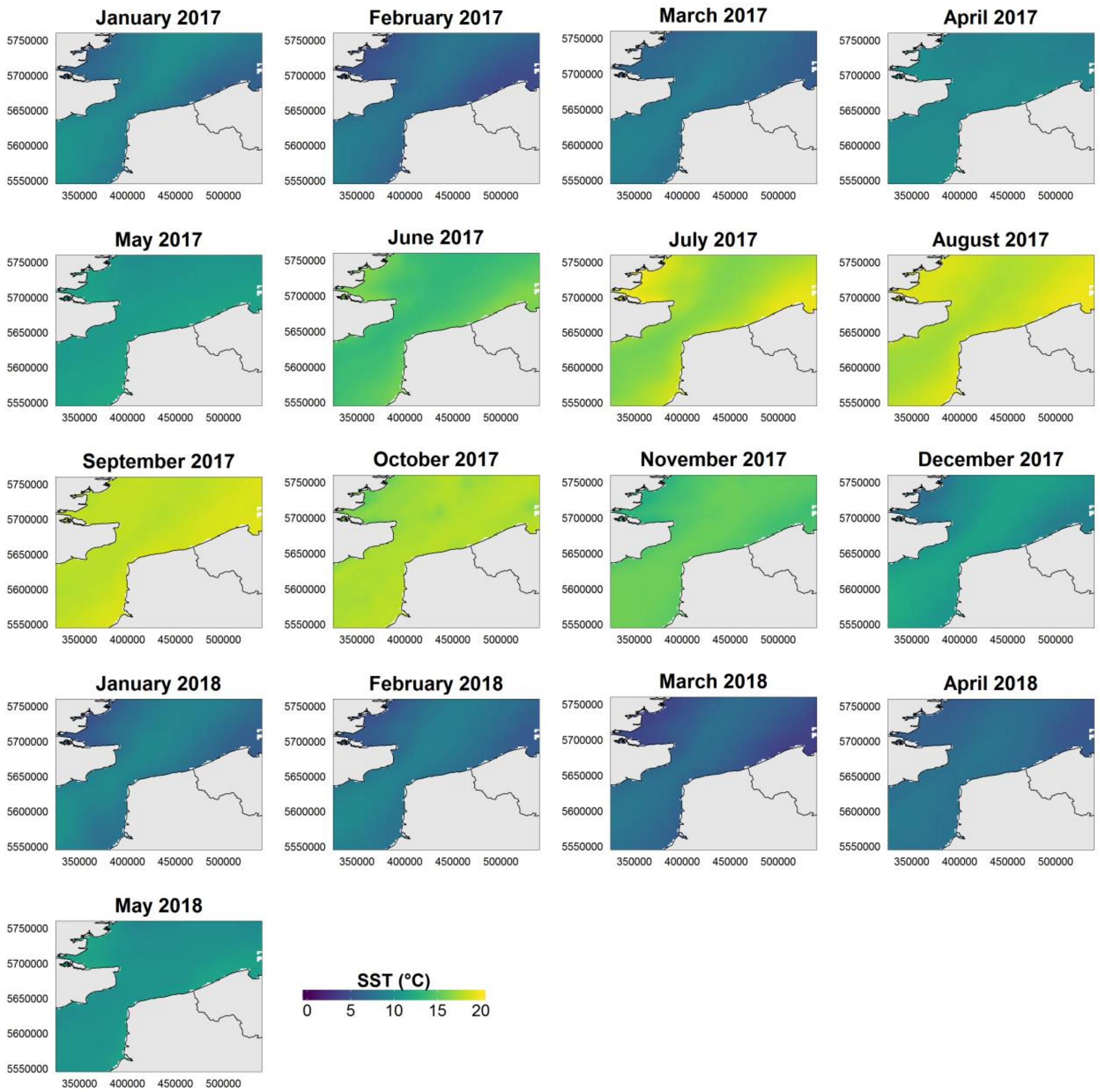


**Figure E.4. Monthly distribution of sea surface temperature (SST).** Average distribution for all months between January 2017 and May 2018. The surface temperature was extracted from NASA data (https://podaac.jpl.nasa.gov/dataset/JPL_OUROCEAN-L4UHfnd-GLOB-G1SST).

### Appendix F. Sightings recorded during each flight session.

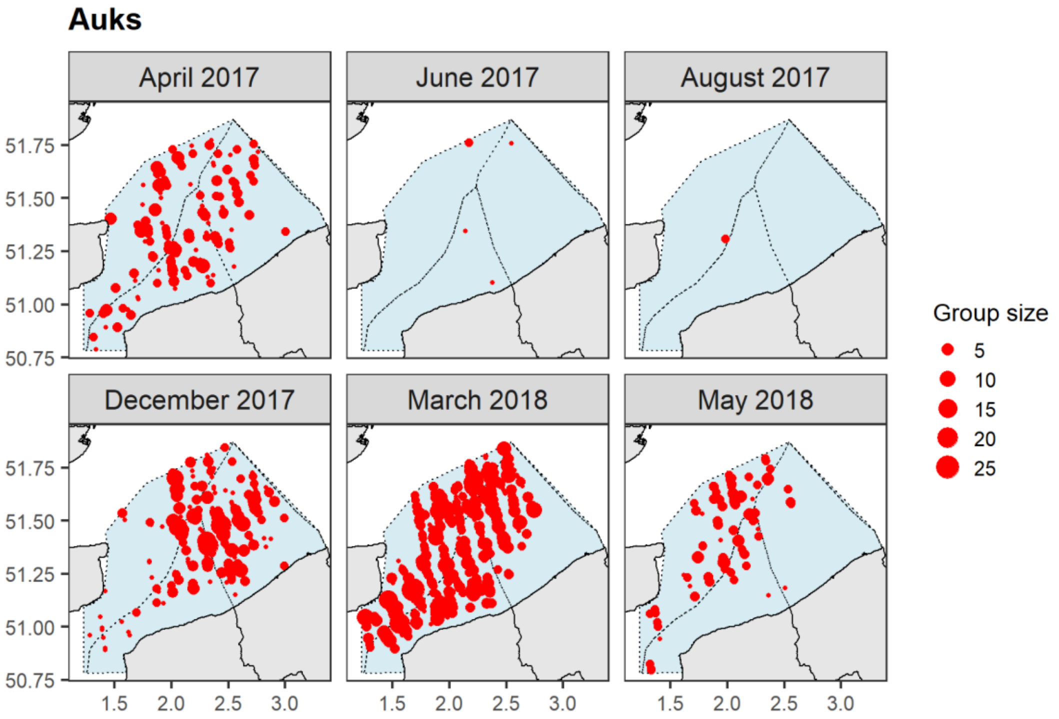


**Figure F.1. Auk sightings.** The red dots represent the sightings and the size of the symbols represents the group size in number of individuals; the light blue dashed area represents the study area.

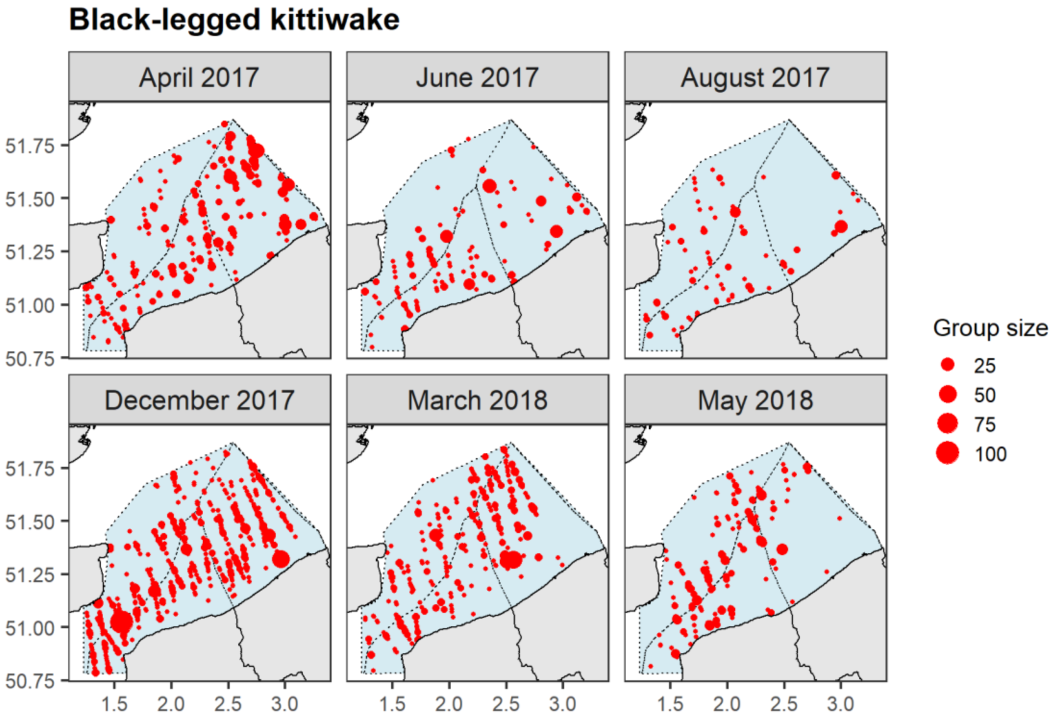


**Figure F.2. Black-legged kittiwake sightings.** The red dots represent the sightings and the size of the symbols represents the group size in number of individuals; the light blue dashed area represents the study area.

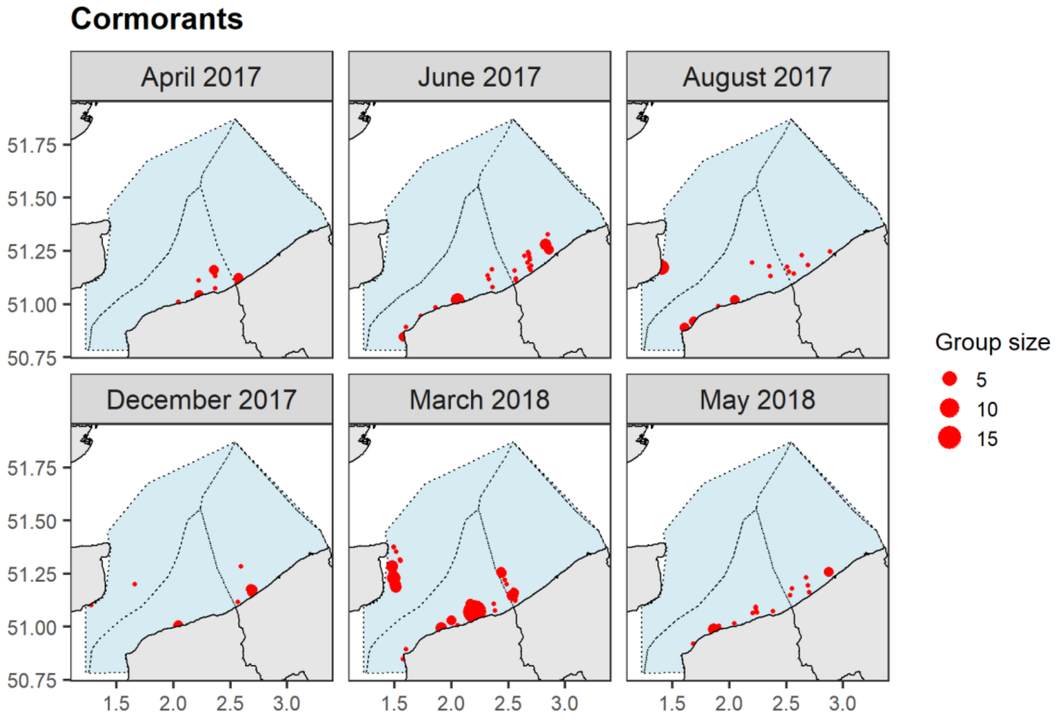


**Figure F.3. Cormorant sightings.** The red dots represent the sightings and the size of the symbols represents the group size in number of individuals; the light blue dashed area represents the study area.

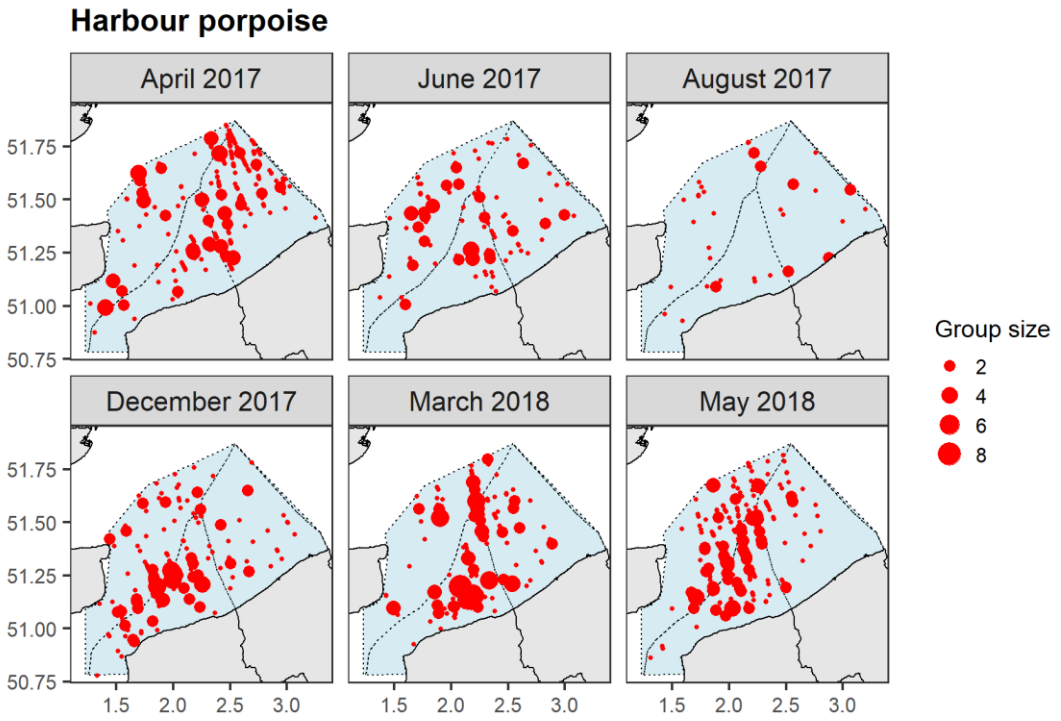


**Figure F.4. Harbour porpoise sightings.** The red dots represent the sightings and the size of the symbols represents the group size in number of individuals; the light blue dashed area represents the study area.

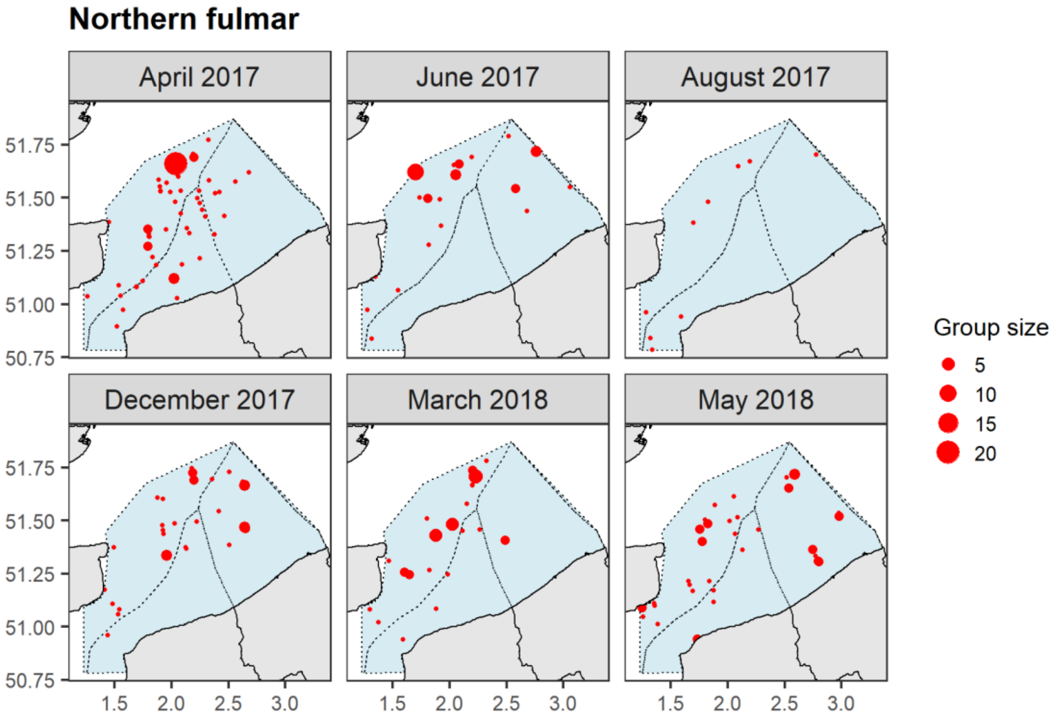


**Figure F.5. Northern fulmar sightings.** The red dots represent the sightings and the size of the symbols represents the group size in number of individuals; the light blue dashed area represents the study area.

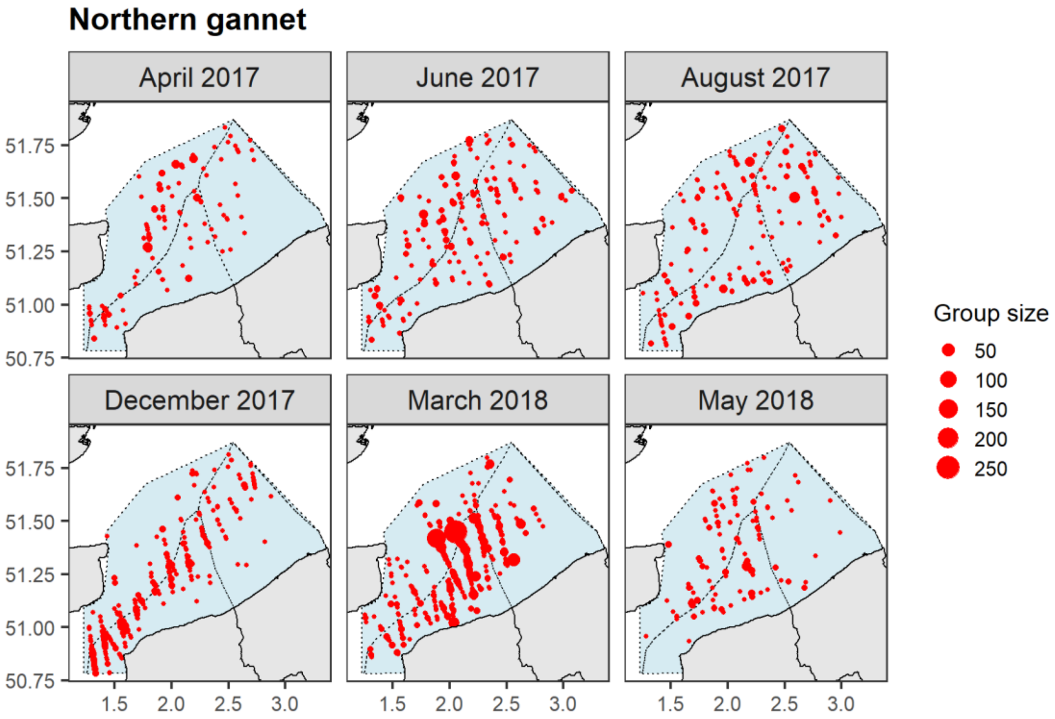


**Figure F.6. Northern gannet sightings.** The red dots represent the sightings and the size of the symbols represents the group size in number of individuals; the light blue dashed area represents the study area.

**Appendix G.** **The monthly functional relationships between species and the selected variables**. Solid lines represent the estimated smooth functions for each month of the study period and the blue shaded regions show the approximate 95% confidence intervals. The density of individuals is shown on the y-axis, where a zero indicates no effect of the covariate. A blank panel indicates that the variable was not selected in the model for this species group. (SST: sea surface temperature; Chl a: chlorophyll a concentration; D. coast: distance to coast; D. wind farms: distance to operational wind farms; D. colony: distance to the colony; B-l.: black-legged; H: harbour; N.: northern.).

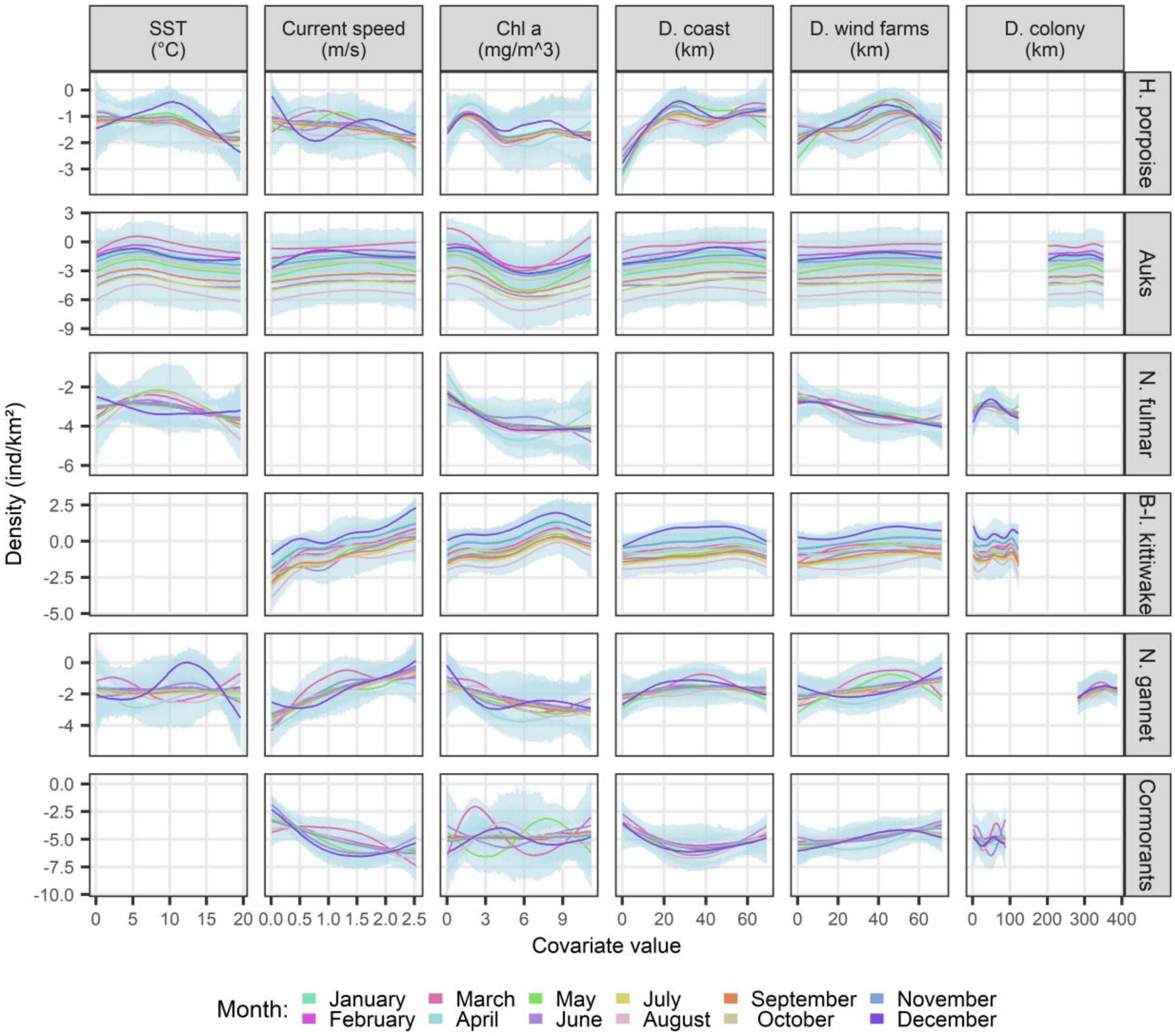

### Appendix H. Monthly predicted densities in the southern North Sea.

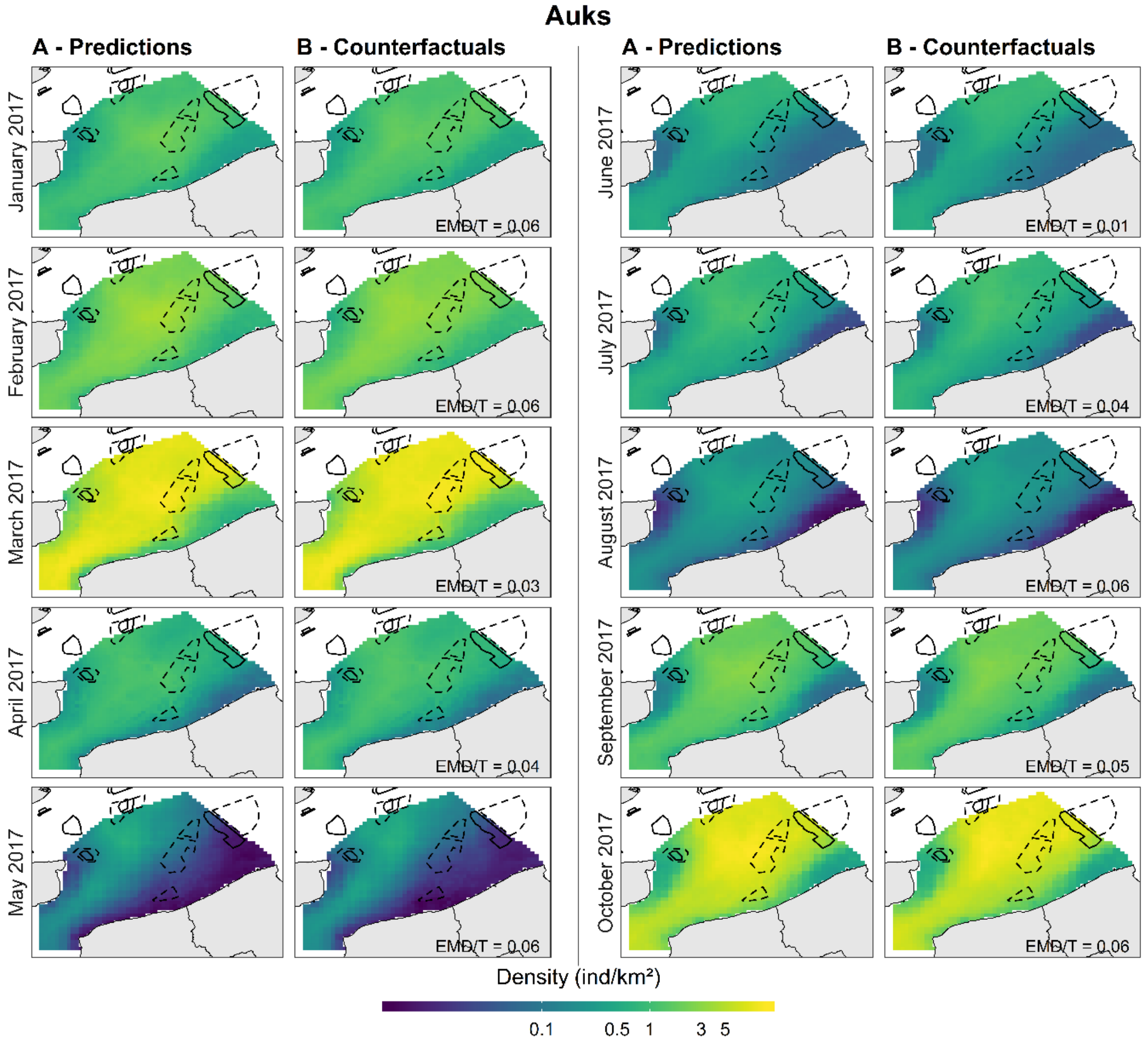


**Figure H.1.1. Monthly predicted densities of auks in the southern North Sea, with model predictions (A) and counterfactuals (B), from January to October 2017**. Black solid boxes represent the wind farms in operation and black dashed boxes the wind farms under development.

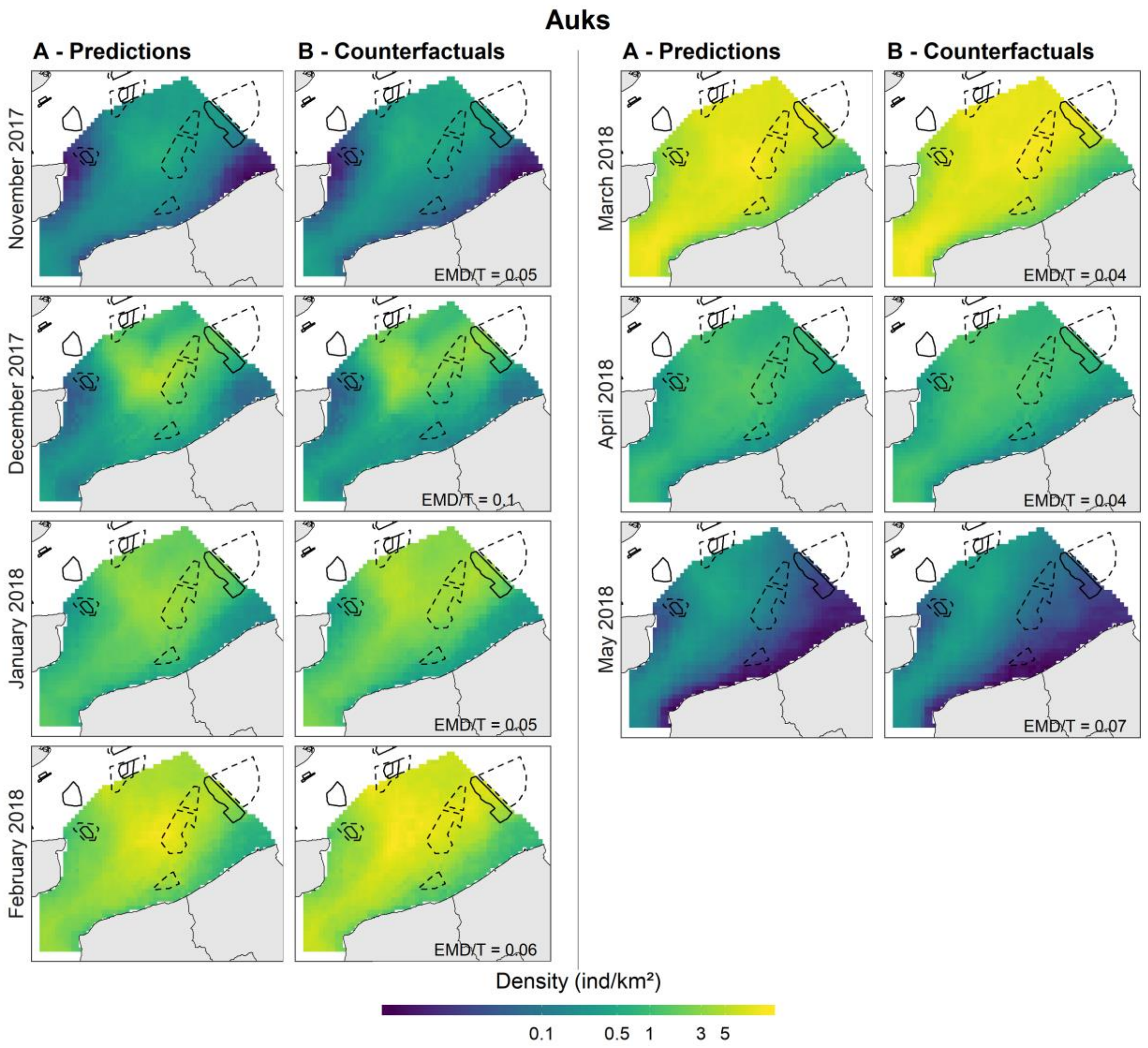


**Figure H.1.2. Monthly predicted densities of auks in the southern North Sea, with model predictions (A) and counterfactuals (B), from November 2017 to May 2018**. Black solid boxes represent the wind farms in operation and black dashed boxes the wind farms under development.

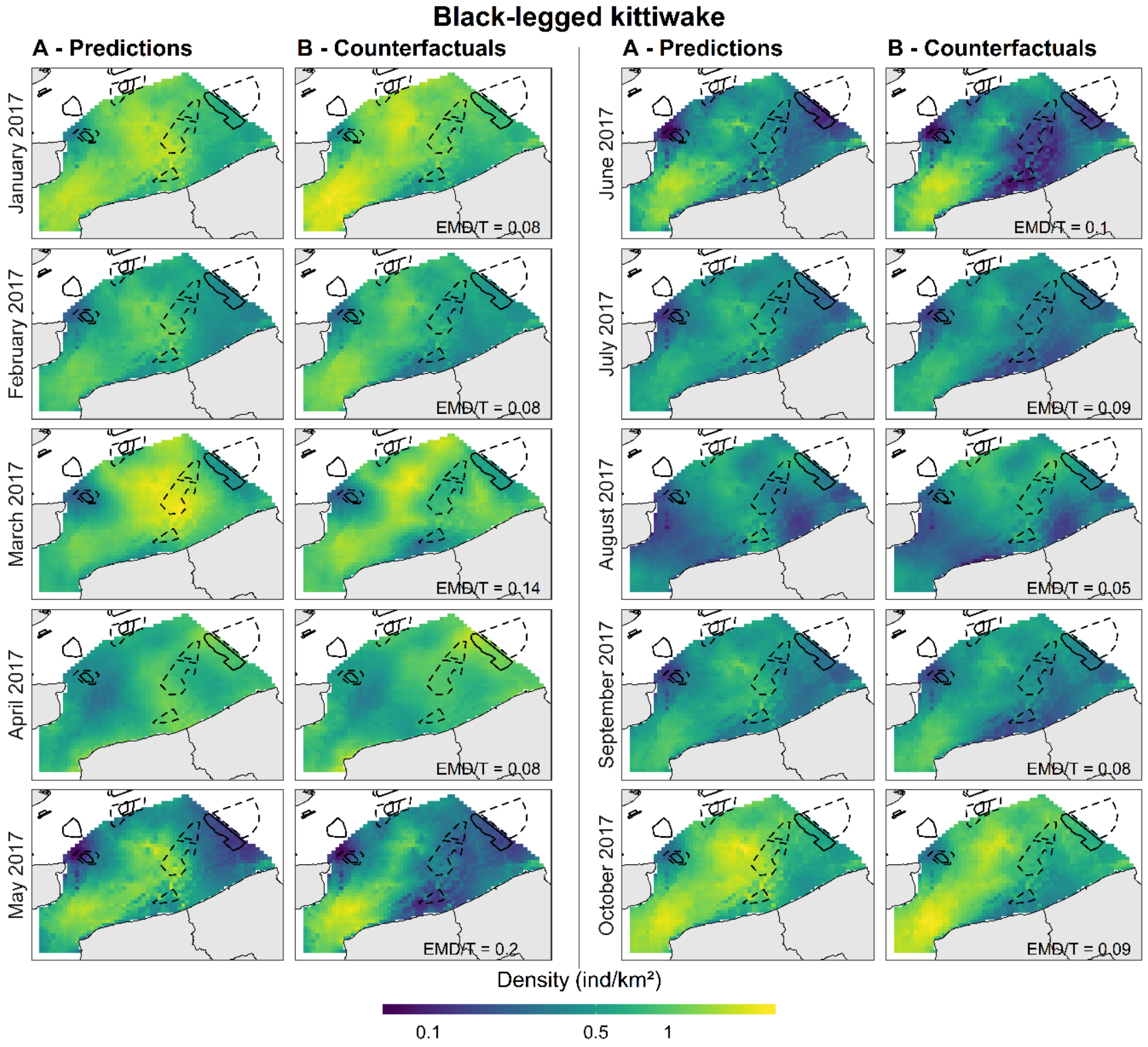


**Figure H.2.1. Monthly predicted densities of black-legged kittiwakes in the southern North Sea, with model predictions (A) and counterfactuals (B), from January to October 2017**. Black solid boxes represent the wind farms in operation and black dashed boxes the wind farms under development.

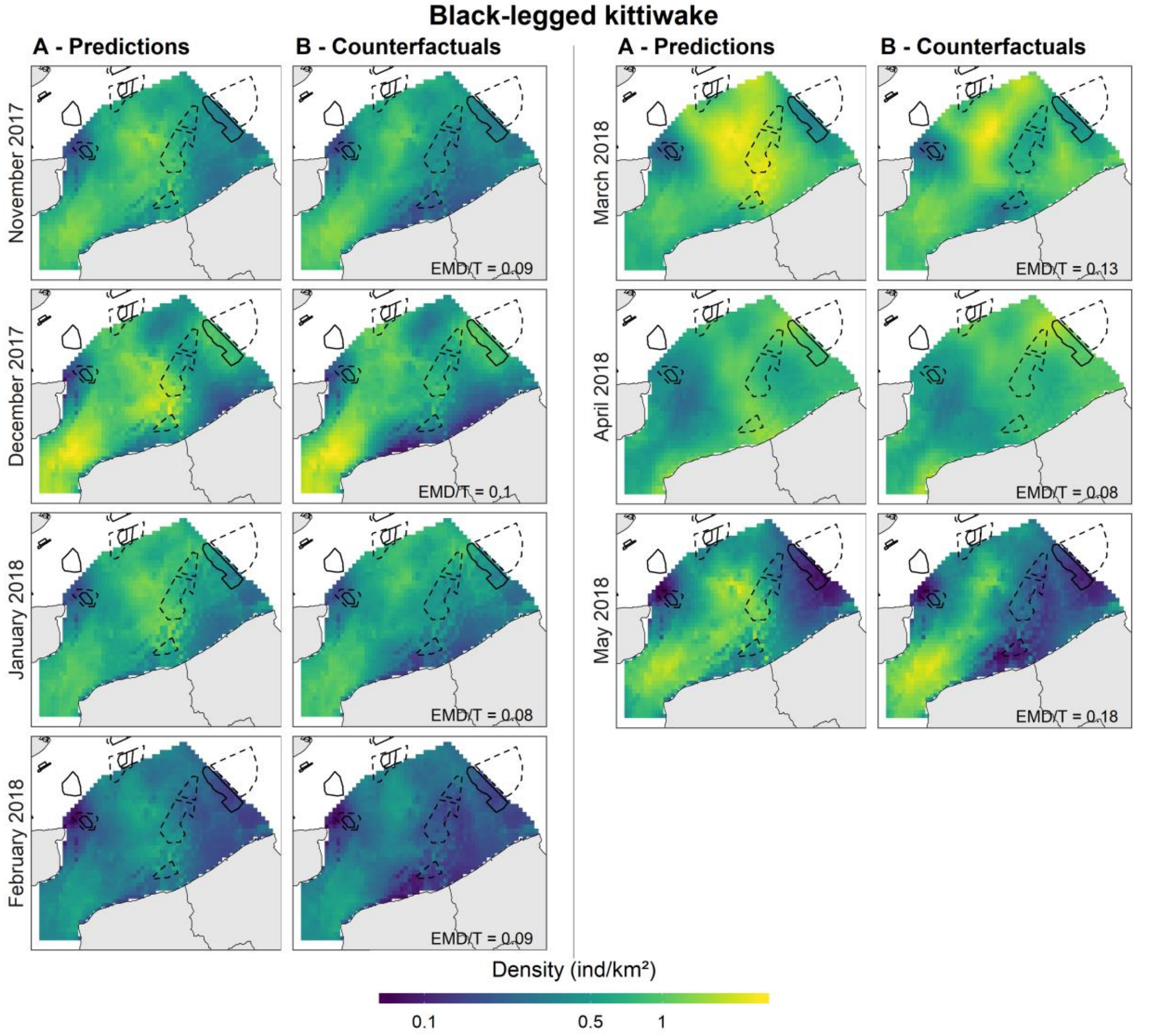


**Figure H.2.2. Monthly predicted densities of auks in the southern North Sea, with model predictions (A) and counterfactuals (B), from November 2017 to May 2018**. Black solid boxes represent the wind farms in operation and black dashed boxes the wind farms under development.

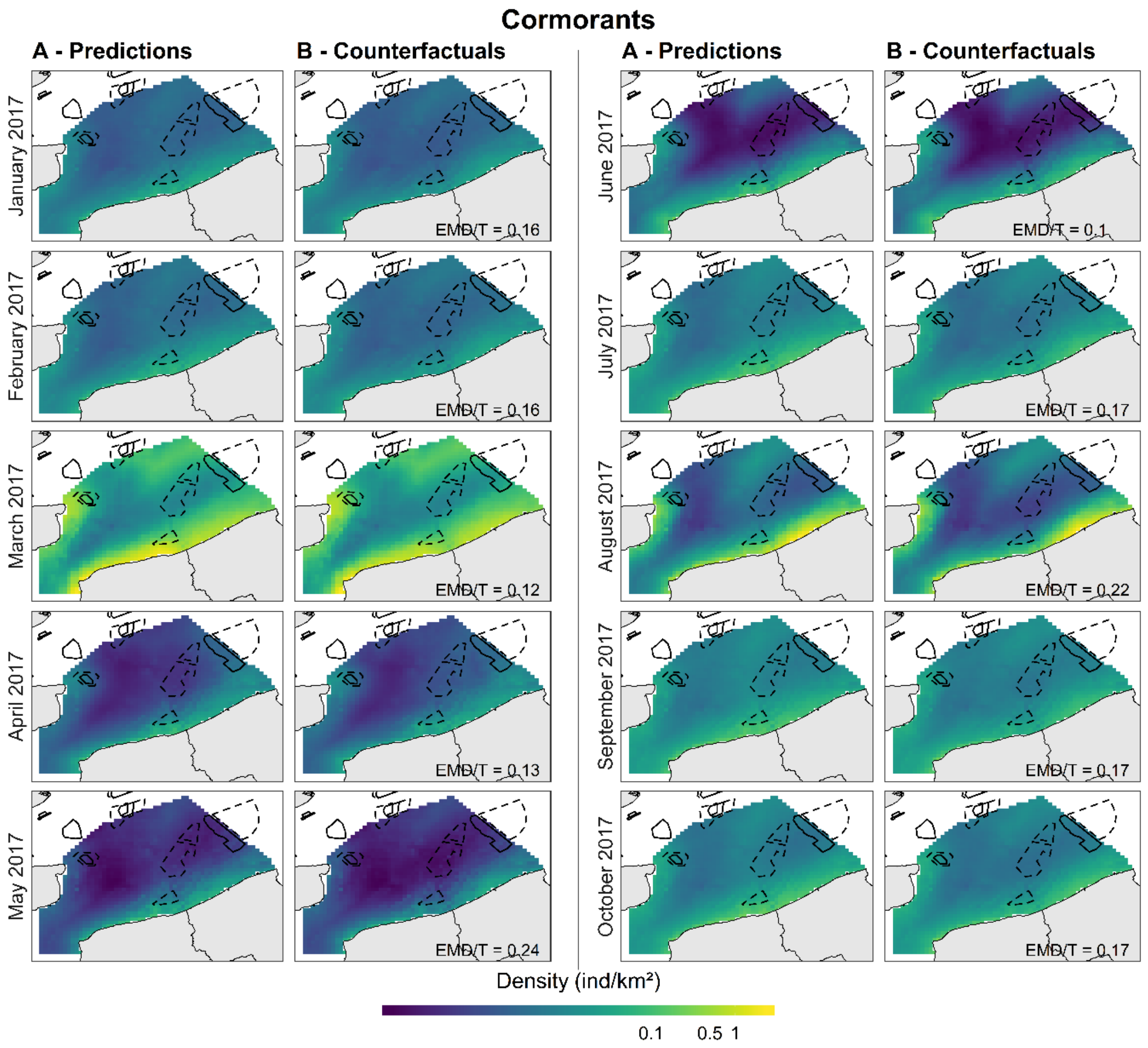


**Figure H.3.1. Monthly predicted densities of cormorants in the southern North Sea, with model predictions (A) and counterfactuals (B), from January to October 2017**. Black solid boxes represent the wind farms in operation and black dashed boxes the wind farms under development.

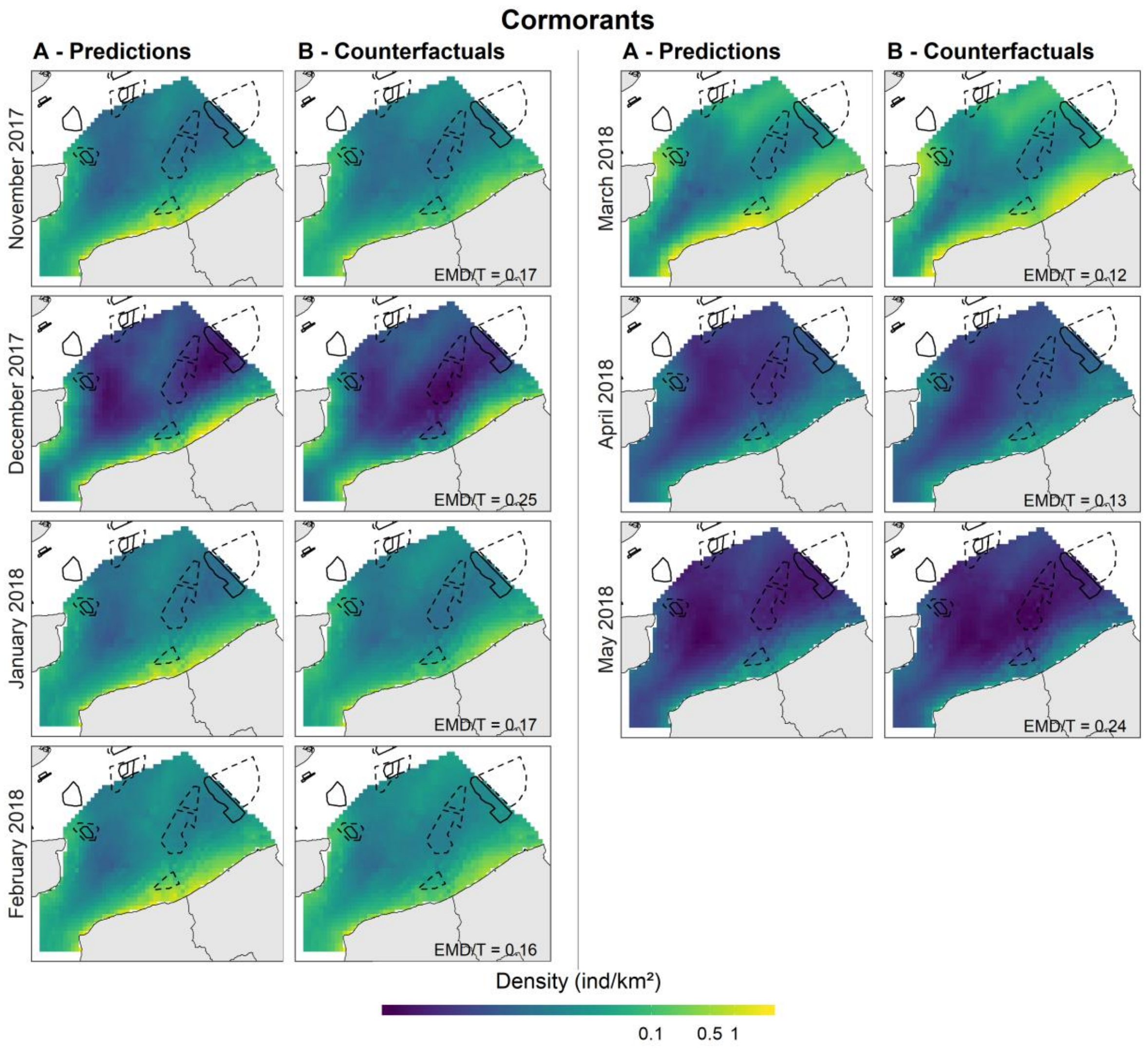


**Figure H.3.2. Monthly predicted densities of auks in the southern North Sea, with model predictions (A) and counterfactuals (B), from November 2017 to May 2018**. Black solid boxes represent the wind farms in operation and black dashed boxes the wind farms under development.

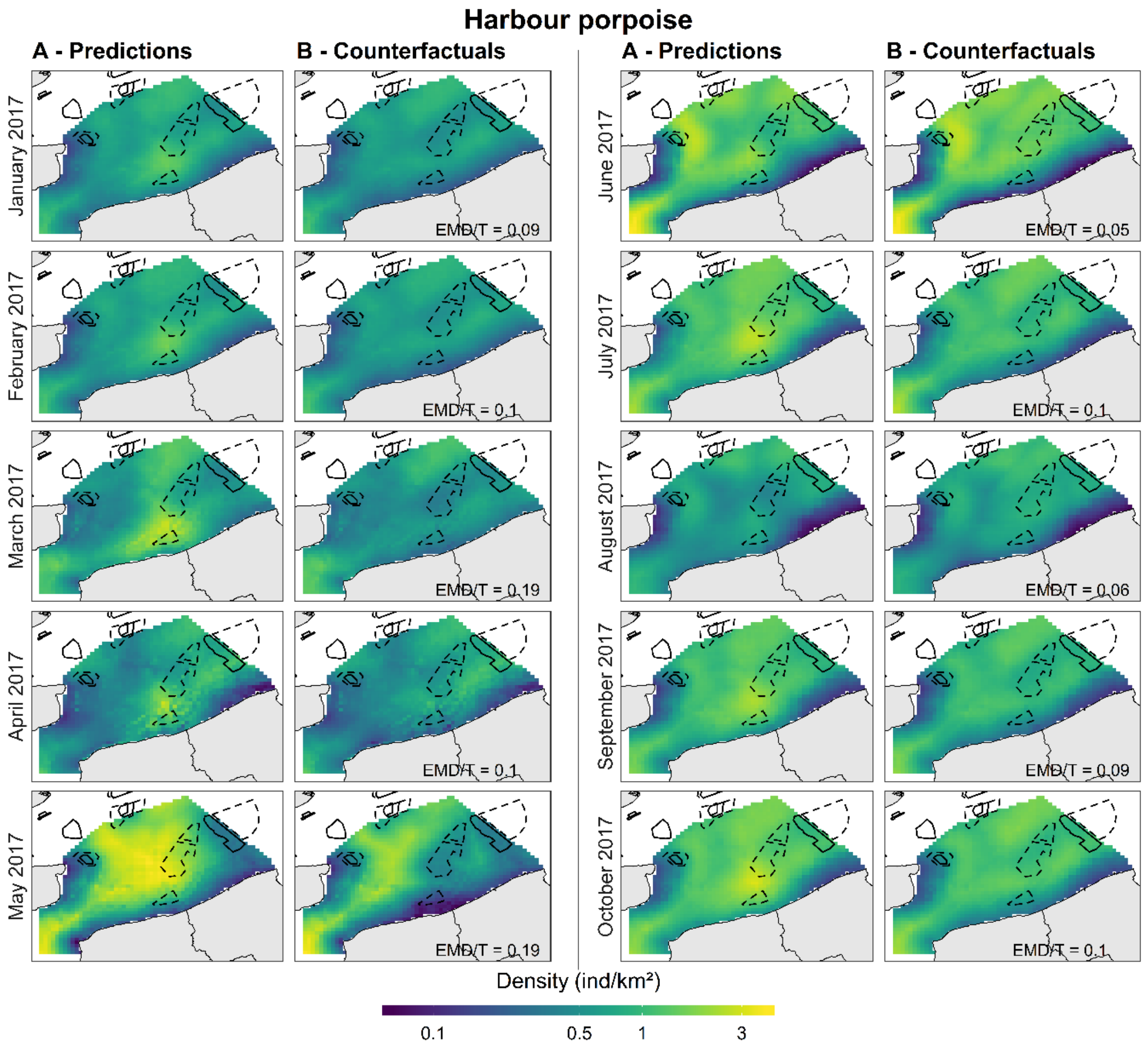


**Figure H.4.1. Monthly predicted densities of harbour porpoises in the southern North Sea, with model predictions (A) and counterfactuals (B), from January to October 2017**. Black solid boxes represent the wind farms in operation and black dashed boxes the wind farms under development.

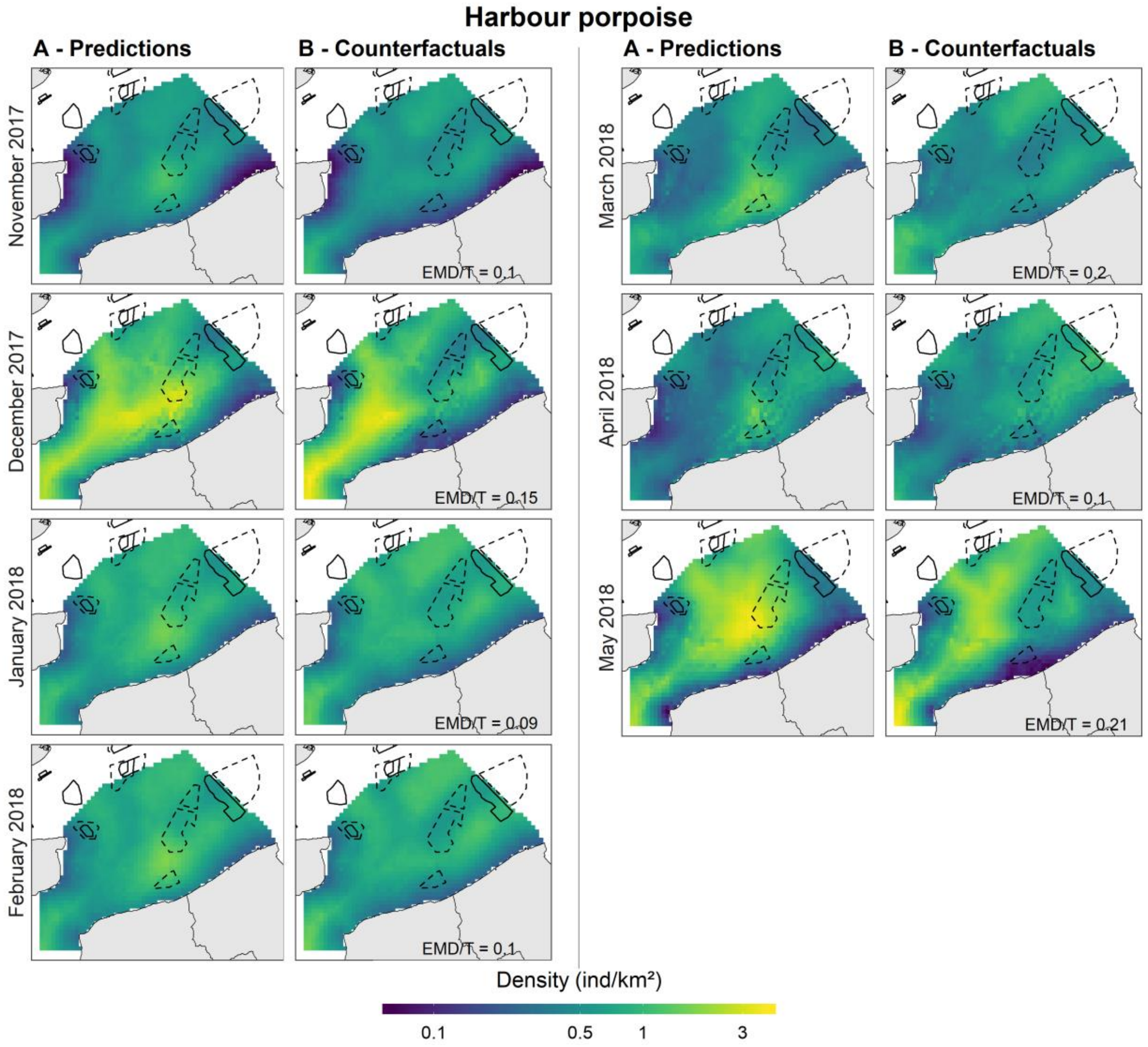


**Figure H.4.2. Monthly predicted densities of auks in the southern North Sea, with model predictions (A) and counterfactuals (B), from November 2017 to May 2018**. Black solid boxes represent the wind farms in operation and black dashed boxes the wind farms under development.

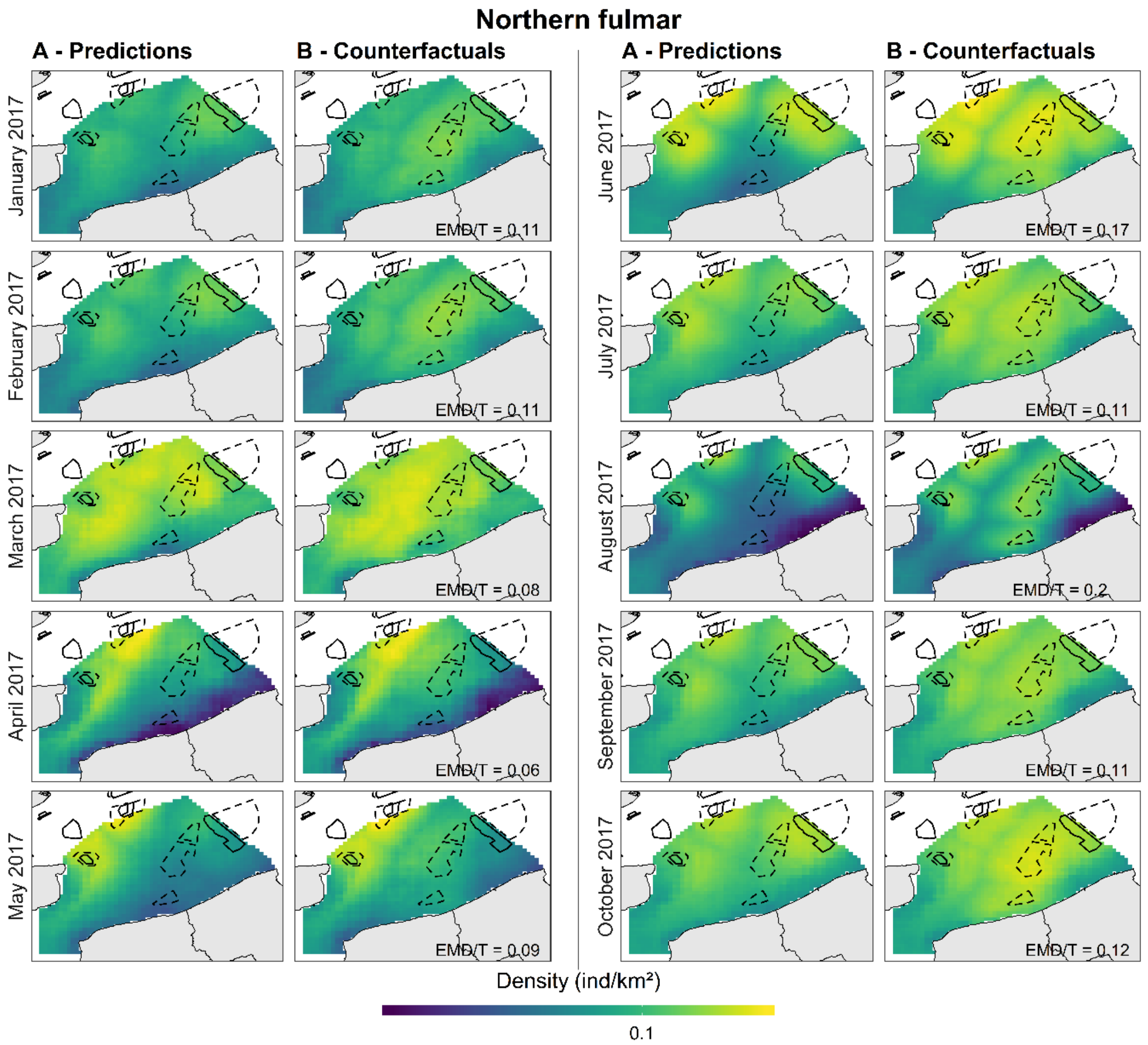


**Figure H.5.1. Monthly predicted densities of northern fulmars in the southern North Sea, with model predictions (A) and counterfactuals (B), from January to October 2017**. Black solid boxes represent the wind farms in operation and black dashed boxes the wind farms under development.

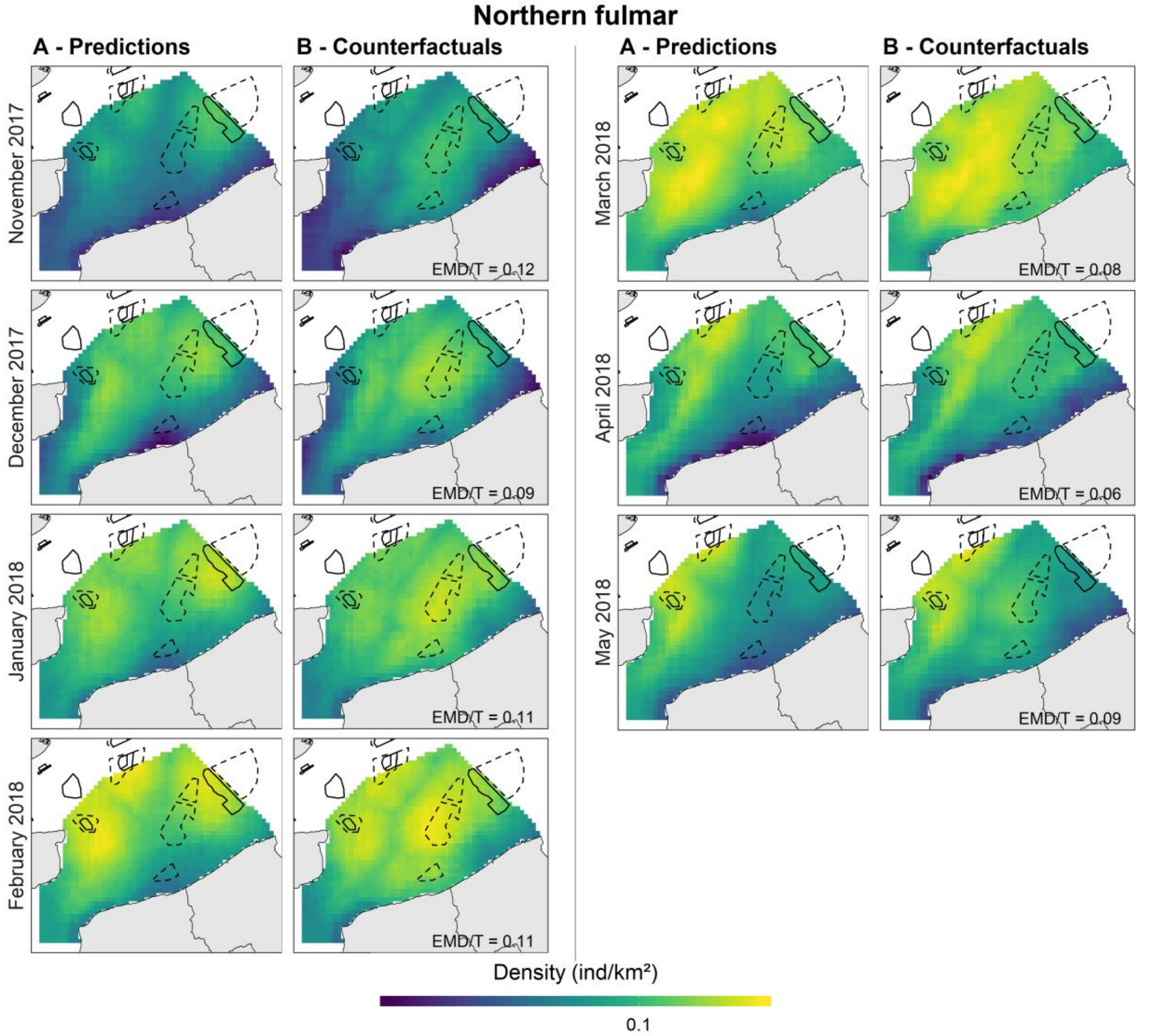


**Figure H.5.2. Monthly predicted densities of northern fulmars in the southern North Sea, with model predictions (A) and counterfactuals (B), from November 2017 to May 2018**. Black solid boxes represent the wind farms in operation and black dashed boxes the wind farms under development.

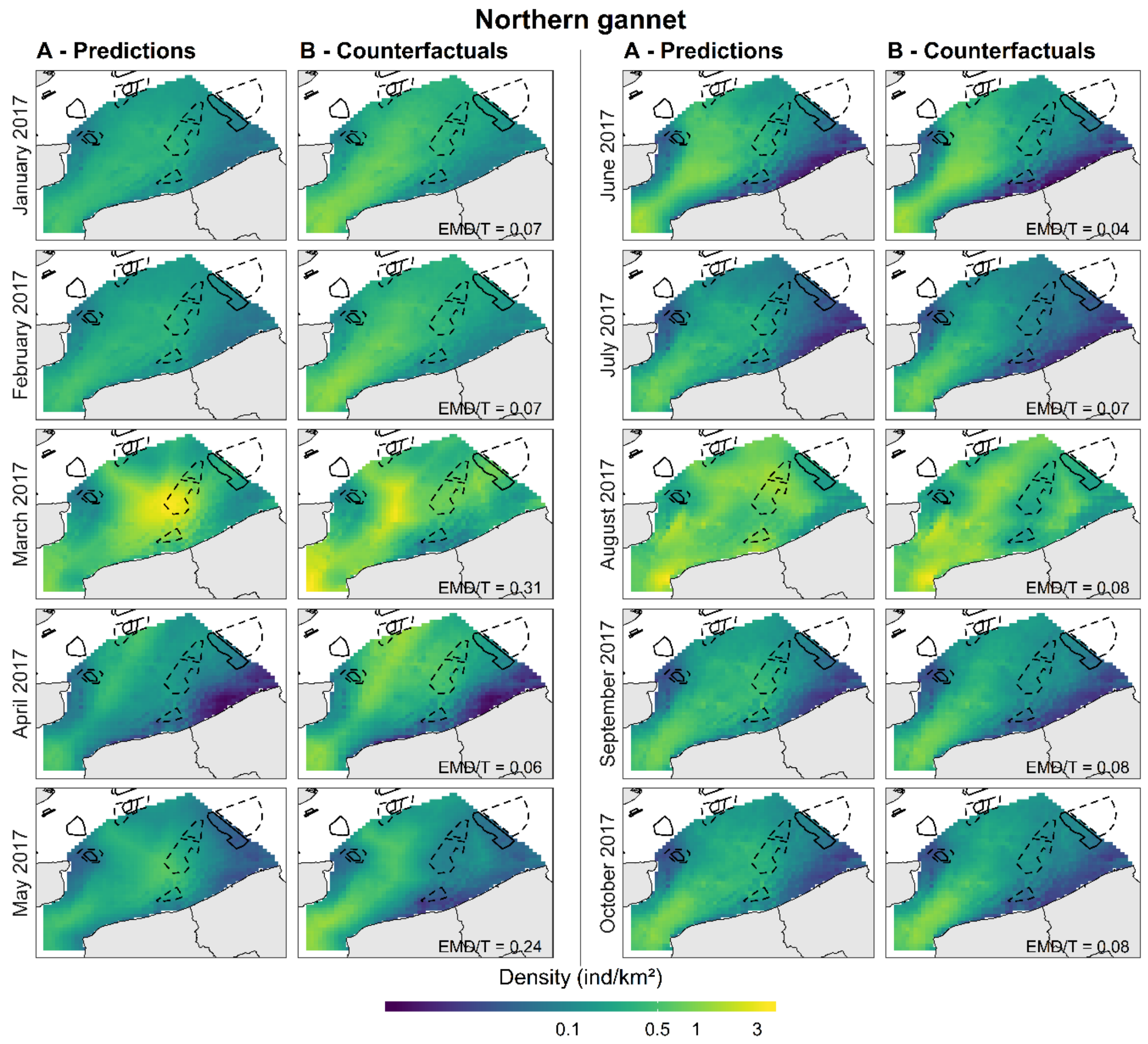


**Figure H.6.1. Monthly predicted densities of northern gannets in the southern North Sea, with model predictions (A) and counterfactuals (B), from January to October 2017**. Black solid boxes represent the wind farms in operation and black dashed boxes the wind farms under development.

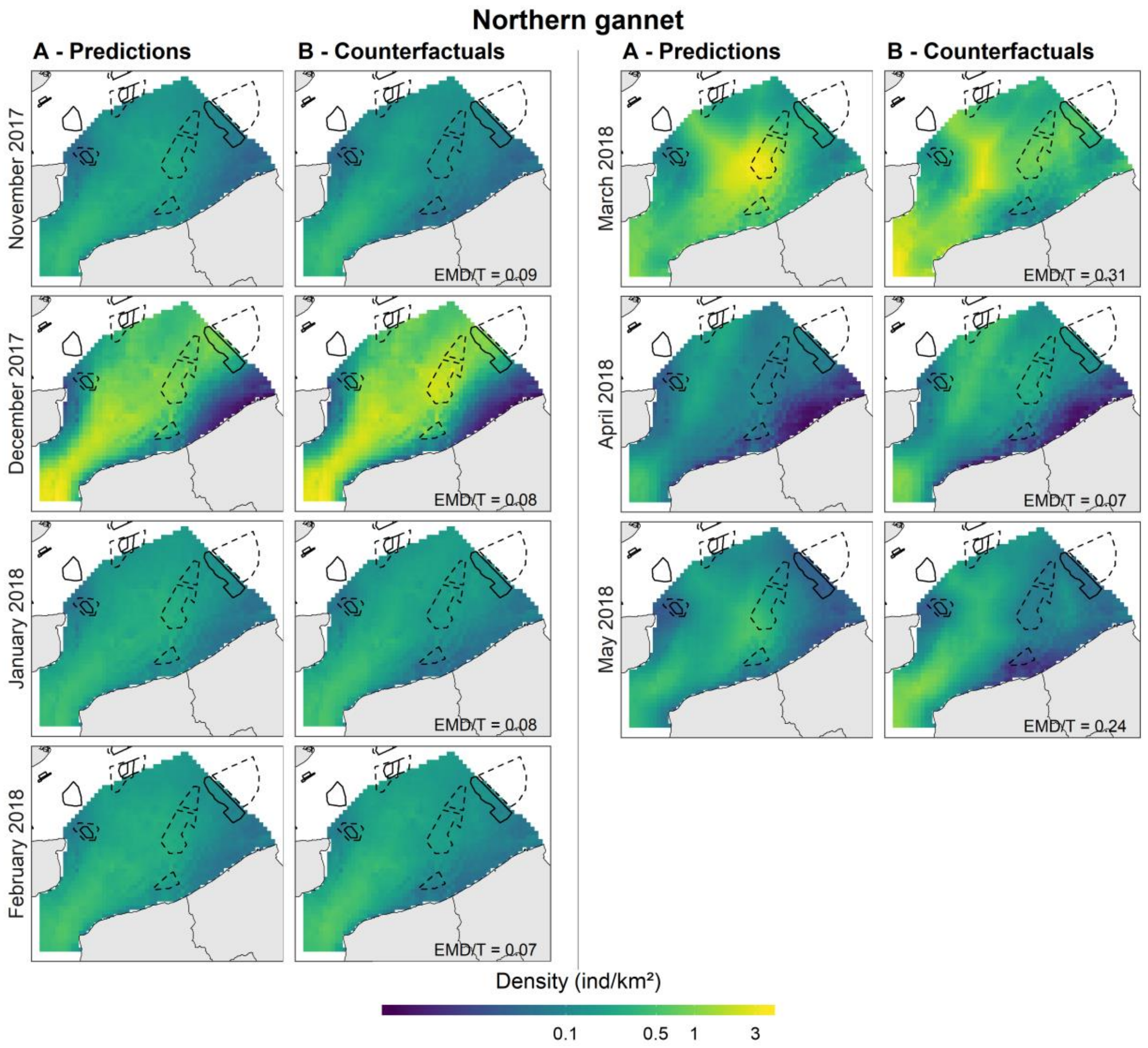


**Figure H.6.2. Monthly predicted densities of northern gannets in the southern North Sea, with model predictions (A) and counterfactuals (B), from November 2017 to May 2018**. Black solid boxes represent the wind farms in operation and black dashed boxes the wind farms under development.

**Appendix I. Monthly abundances in number of individuals estimated from the model (A) and the counterfactuals (B).** Abundance estimates are shown for each month between January 2017 and May 2018. Grey shaded regions represent confidence intervals. Confidence intervals are very tight around the abundance estimates when a flight session is associated with that month, and are much wider when no flight session happened that month (the uncertainty is greater), particularly between September and November because the models were calibrated on the flight sessions in August and December.

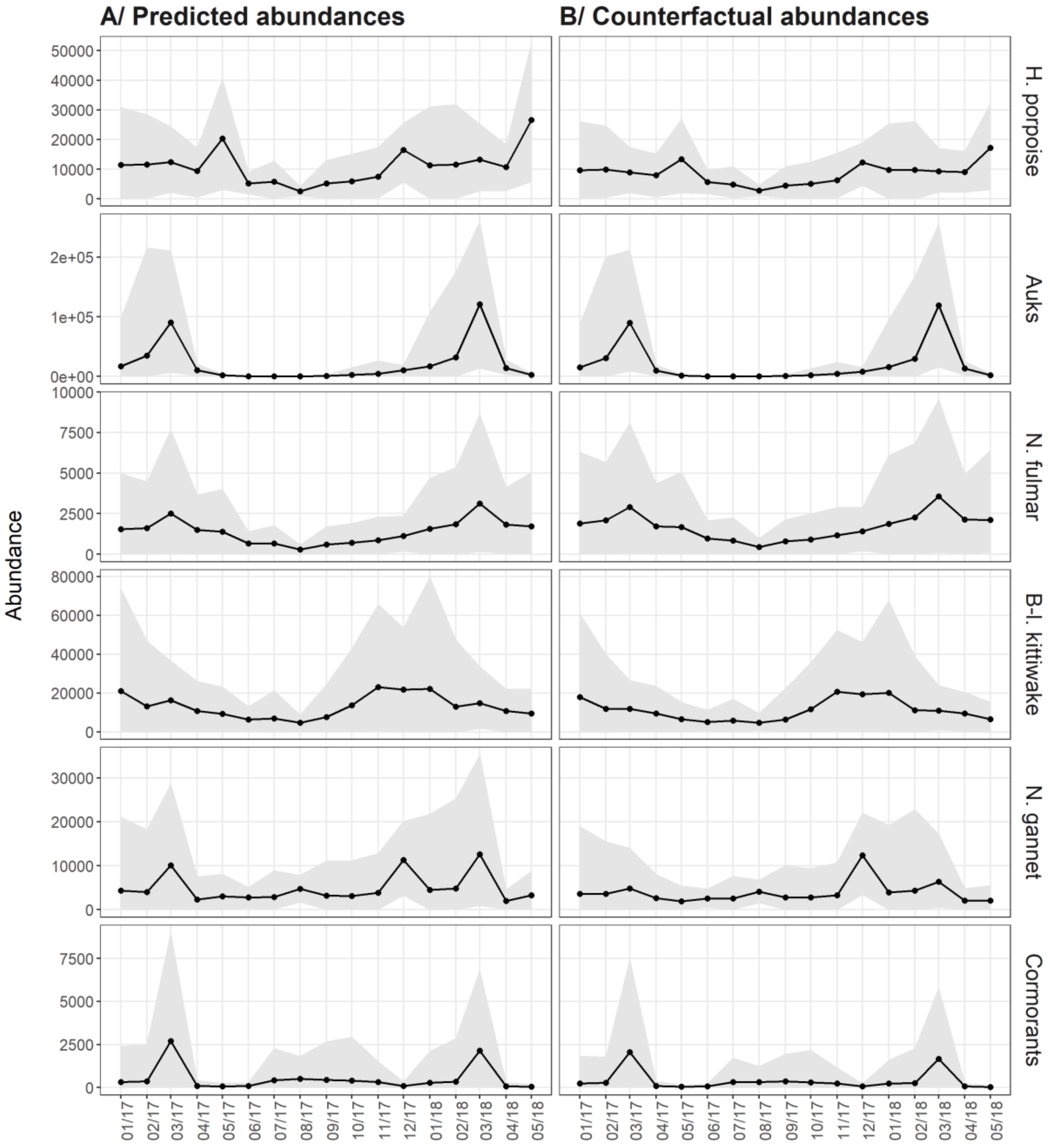
